# An integrated platform for genome-wide functional analysis in *Aspergillus niger* through *Ac/Ds* insertion profiling and metabolic modeling

**DOI:** 10.64898/2026.09.24.754008

**Authors:** Kangzhou Huang, Yikun Lin, Bin Wang, Li Pan

**Author notes:** Contributing authors (Kangzhou Huang, Yikun Lin, Bin Wang).

## Abstract

Genome-scale metabolic models guide rational strain engineering, while high-throughput insertional mutagenesis enables systematic investigation of gene function and condition-dependent growth requirements. However, a platform integrating high-density transposon insertion sequencing with matched stoichiometric, thermodynamic, enzyme-constrained and combined metabolic models has not, to our knowledge, been reported in *Aspergillus*. Here, we establish a unified experimental and computational platform in *Aspergillus niger* ATCC 1015 by combining curated metabolic reconstruction, *Ac/Ds* transposon mutagenesis and machine-learning-based essentiality prediction. Our stoichiometric reconstruction, FBA1015, expands reaction coverage and compartmental resolution beyond four published reconstructions. From FBA1015, we developed three models sharing its biochemical network: GECKO light with enzyme-capacity constraints, TFA with thermodynamic constraints and ecTFA combining both. FBA1015 and its derivatives performed competitively in phenotype benchmarks, with added constraints improving agreement with measured growth and gas exchange. Insertion libraries grown in rich medium, glucose or xylose achieved genome-wide densities of one distinct site per 21–112 bp across individual libraries. A machine-learning classifier integrated insertion patterns and gene structure to predict essentiality across 11,111 genes. Applying a stringent consensus criterion identified 709 genes predicted essential on both glucose and xylose as sole carbon sources, and 91 predicted essential on one but nonessential on the other. Integration with simulated gene deletions prioritizes candidate genes and testable pathway hypotheses, connecting high-throughput genetic screening with mechanistic analysis for fungal functional investigation and strain engineering.

## 1 Introduction

*Aspergillus niger* is an established producer of organic acids and industrial enzymes. Genome-scale metabolic models (GEMs) support its rational engineering by connecting genes and reactions to growth and production. Successive reconstructions have expanded the species’ metabolic representation through updated annotations, community curation and enzyme constraints [1–6]. Thermodynamic constraints restrict feasible reaction directions, while enzyme constraints limit catalytic capacity; both can supplement stoichiometric flux balance analysis [7–9]. Applying these constraints to a common network allows their effects to be distinguished from differences in reconstruction content. Experimental perturbation data provide a complementary test of the gene requirements represented by that network.

Genome-wide perturbation resources have transformed fungal functional genetics. Systematic deletion collections enabled large-scale phenotyping in *Saccharomyces cerevisiae* and *Neurospora crassa* [10–12], and pooled CRISPR–Cas9 libraries now support single and combinatorial gene inactivation in *A. niger* [13]. A randomly barcoded insertion library has also enabled pooled fitness measurements in *Trichoderma atroviride* [14]. Transposon insertion sequencing (Tn-seq) offers a complementary approach by recovering insertion patterns across the genome. Saturated transposition can resolve gene-level and subgenic effects [15], while supervised analysis of insertion features has supported essentiality prediction in *Candida albicans* and *Cryptococcus neoformans* [16, 17].

Interpreting these patterns in filamentous fungi requires accounting for both insertion opportunity and biological selection. Genetically distinct nuclei can share a cytoplasm, allowing essential-gene disruptions to persist through heterokaryotic complementation [18]. Insertions may also preserve function depending on their position within a gene [15, 17], whereas sparse recovery can reflect poor chromatin accessibility. Published ATAC-seq maps provide an independent reference for assessing the latter association [19]. Consequently, insertion presence or absence alone need not establish whether a gene is required for growth.

Here, we integrate metabolic modelling and high-density *Ac/Ds* insertion profiling in *A. niger* ATCC 1015. FBA1015 and three constrained models derived from its curated network provide a common basis for phenotype, flux and growth comparisons. Complementary insertion libraries define genome-wide patterns under rich-medium, glucose and xylose conditions. A machine-learning classifier combines insertion patterns with gene structure, and a stringent consensus criterion selects stable predictions for model comparison. Together, these resources identify shared and carbon-source-associated essentiality predictions and connect individual disagreements to testable pathway assumptions.

## 2 Results

### 2.1 Reconstruction of FBA1015 and derivation of constrained models

FBA1015 provides a common metabolic network for comparing four models with different constraints (Fig. 1A). Homology-derived drafts were integrated with a compartment-expanded iJB1325 reconstruction and manually curated. The resulting network formed the basis of stoichiometric FBA, thermodynamic flux analysis (TFA), enzyme-constrained GECKO light and combined enzyme-constrained TFA (ecTFA), allowing the effects of added constraints to be assessed on the same reconstruction.

**Fig. 1.**
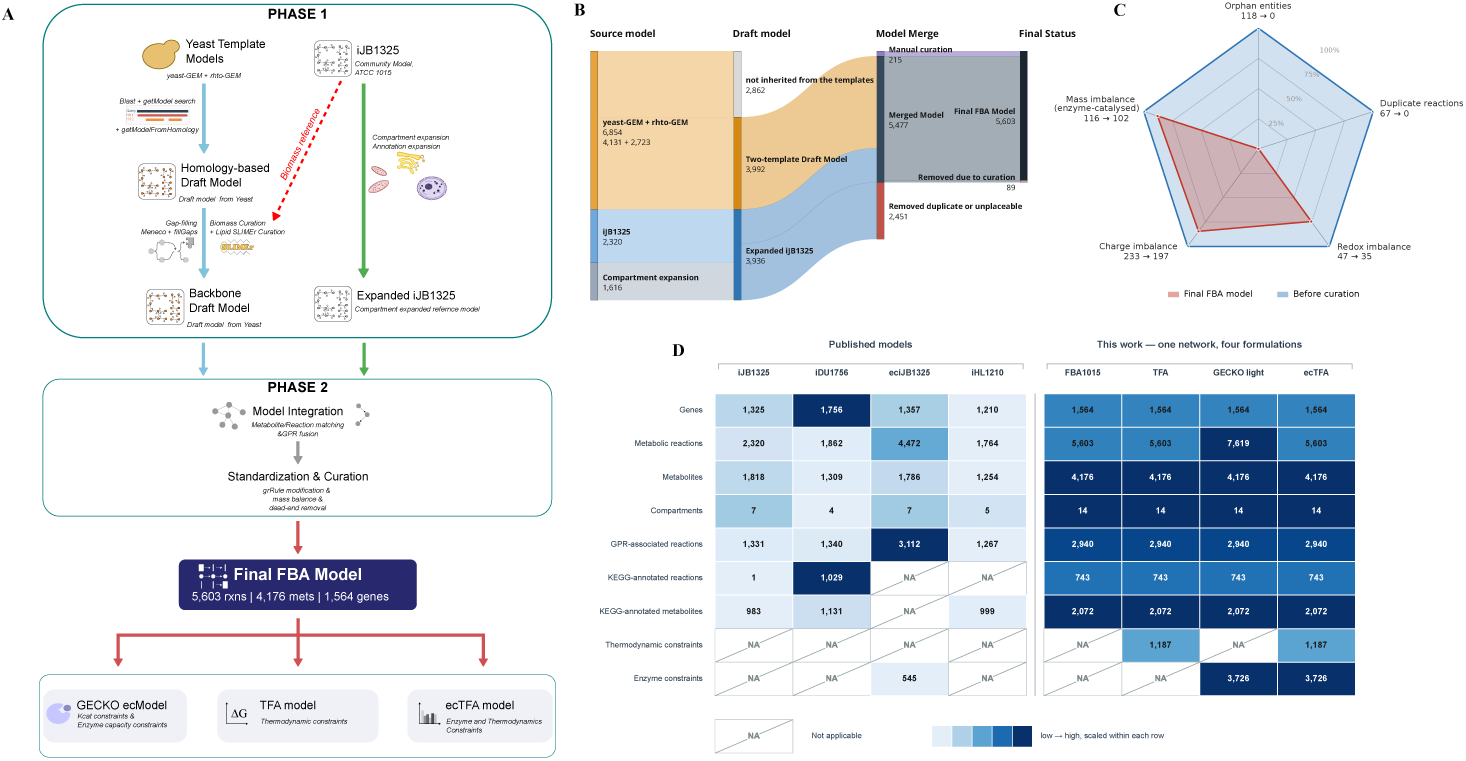
Reconstruction of FBA1015 and its constrained models. (**A**) Reconstruction workflow: homology-derived drafts and compartment-expanded iJB1325 were merged and curated, then used to derive three constrained models. (**B**) Reaction provenance and retention during reconstruction; manual additions are shown separately from template-derived reactions. (**C**) Network defects before (blue outline) and after (red) curation. Counts are scaled to the larger value within each category; smaller radii indicate fewer defects. (**D**) Model content and annotation coverage, with colour scaled within each row. Directed reaction counts in enzyme-constrained models include the splitting of reversible reactions.

Network integration consolidated overlapping template content and incorporated targeted additions (Fig. 1B). Merging the drafts yielded 5,477 reactions; subsequent curation retired 89 reactions and introduced 215. The final reconstruction therefore combines transferred metabolic knowledge with manual additions rather than simply accumulating reactions from its source models.

Curation reduced all five assessed classes of network defects (Fig. 1C). Orphan entities (unconnected metabolites and genes) and duplicate reaction pairs were eliminated, while the counts of redox-, charge- and mass-imbalanced reactions decreased. The reductions improved network consistency without eliminating all biochemical imbalances.

FBA1015 expands reaction coverage and compartmental resolution relative to the four published reconstructions (Fig. 1D). It contains 5,603 metabolic reactions and 4,176 metabolites across 14 compartments, compared with 1,764–4,472 reactions and four to seven compartments in the published models. Annotation coverage varied by metric: FBA1015 contained the most annotated metabolites, whereas published models retained larger gene sets or more reactions with gene rules or KEGG annotations. The larger reaction counts of enzyme-constrained models partly reflect the splitting of reversible reactions, rather than additional metabolic chemistry.

### 2.2 Model performance across phenotype, flux and growth benchmarks

FBA1015 and its constrained models combined broad phenotype coverage with improved prediction after adding enzyme constraints (Fig. 2A). The iJB1325 benchmark tests growth, biomass-component synthesis and specified reaction-flux outcomes. Three of its 471 cases specified outcomes incompatible with their own reaction bounds and were excluded for every model. All four models developed here were evaluable on the same 428 of the remaining 468 cases (91.5%). GECKO light and ecTFA increased MCC from 0.432 for FBA1015 to 0.494 and 0.497, respectively, approaching the highest point estimate of 0.516 for eciJB1325. The shared case set supports direct comparison within our model family; comparisons with published models use their respective evaluable subsets.

**Fig. 2.**
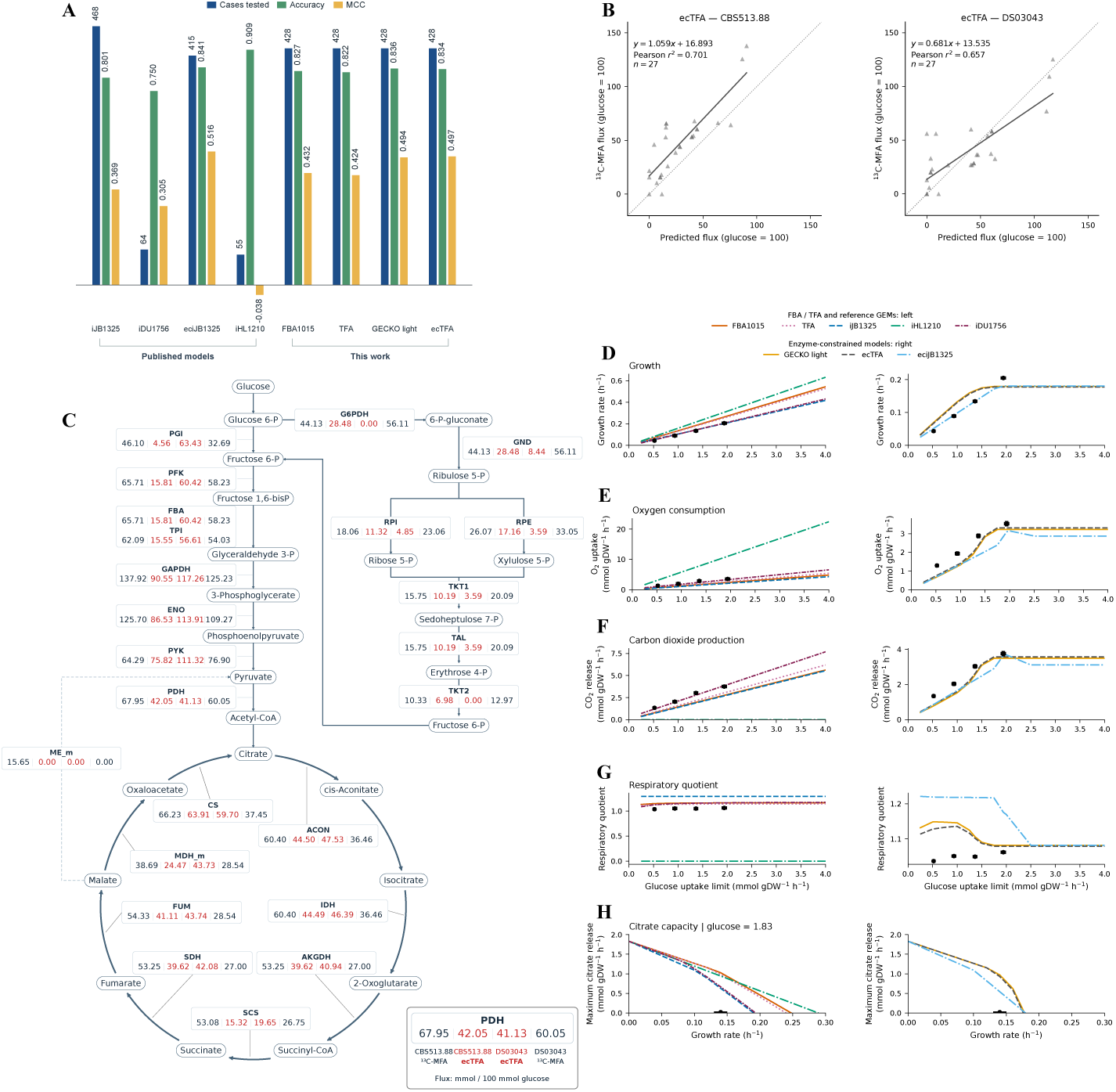
Model performance across phenotype, flux and growth benchmarks. (**A**) Coverage, accuracy and Matthews correlation coefficient (MCC) for the iJB1325 benchmark of growth, biomass-component synthesis and specified reaction-flux outcomes [4]. Of 471 original cases, three specified outcomes incompatible with their own reaction bounds and were excluded for every model, leaving 468 eligible cases. Blue bars indicate evaluable-case counts; accuracy and MCC use the corresponding fractional scale. Metrics are calculated on each model’s evaluable subset. (**B**) Measured and ecTFA-predicted ^13^C-MFA fluxes for two strains [20], using 27 mapped carbon transformations. Dotted line, *y* = *x*; solid line, ordinary least-squares fit. (**C**) Central-carbon fluxes in the order CBS 513.88 measured, CBS 513.88 ecTFA, DS03043 ecTFA and DS03043 measured, in mmol per 100 mmol glucose. (**D**–**G**) Growth, O_2_ uptake, CO_2_ release and respiratory quotient across glucose-uptake ceilings. Gas fluxes are in mmol gDW*^−^*^1^ h*^−^*^1^; black points are four observations from aerobic, glucose-limited chemostats of *A. niger* NW185 [21]. (**H**) Maximum citrate secretion at prescribed growth rates and fixed glucose uptake of 1.83 mmol gDW*^−^*^1^ h*^−^*^1^; the black point marks measured growth and citrate secretion in a batch culture of the glucoamylase-producing strain DS03043 [20], providing a physiological reference rather than an optimized citrate-production condition. In D–H, the left column shows stoichiometric and published models and the right column enzyme-constrained models.

All four models developed here accommodated the measured growth and exchange rates of both CBS 513.88 and the glucoamylase-producing strain DS03043, enabling comparison with 27 measured intracellular carbon transformations (Fig. 2B; Table 1) [20]. In CBS 513.88, each additional constraint scheme improved flux correlation over FBA1015, with ecTFA achieving the highest *r*^2^ among feasible models (0.701, compared with 0.630 for FBA1015). In DS03043, FBA1015 gave the strongest correlation within our model family (*r*^2^ = 0.790), exceeding iJB1325 (0.760), while eciJB1325 gave the highest overall value (0.860). These comparisons establish useful intracellular-flux prediction across two strain backgrounds and show which constraints improve agreement in each.

**Table 1.** Agreement between predicted and measured ^13^C-MFA fluxes for 27 mapped carbon transformations in two strains [20]. Fluxes and regression intercepts are expressed in mmol per 100 mmol glucose. Fits use measured = slope × predicted + intercept; *r*^2^ is squared Pearson correlation. *Infeasible* indicates that the fixed growth and exchange rates could not be satisfied simultaneously (Methods 3.3).

| Model | CBS 513.88 |  |  | DS03043 |  |  |
| --- | --- | --- | --- | --- | --- | --- |
| | slope | intercept | $r^2$ | slope | intercept | $r^2$ |
| iJB1325 | 0.703 | 20.41 | 0.491 | 0.829 | 13.21 | 0.760 |
| iDU1756 |  | *infeasible* |  |  | *infeasible* |  |
| eciJB1325 |  | *infeasible* |  | 0.934 | 9.77 | 0.860 |
| iHL1210 | 0.762 | 15.80 | 0.664 | 1.163 | 5.17 | 0.578 |
| FBA1015 | 0.730 | 20.33 | 0.630 | 1.101 | 8.15 | 0.790 |
| TFA | 0.737 | 17.61 | 0.680 | 1.064 | 6.26 | 0.784 |
| GECKO light | 1.017 | 17.08 | 0.693 | 0.749 | 12.53 | 0.722 |
| ecTFA | 1.059 | 16.89 | 0.701 | 0.681 | 13.53 | 0.657 |

The pathway map connects this predictive performance to individual central-carbon reactions (Fig. 2C). ecTFA closely reproduced citrate-synthase flux in CBS 513.88 and several glycolytic fluxes in DS03043, including phosphofructokinase and enolase. The larger differences were concentrated in upper glycolysis in CBS 513.88 and in glucose-6-phosphate partitioning and the oxidative pentose phosphate pathway in DS03043. The model also permits glucose oxidation to gluconate, providing an alternative entry route downstream of glucose-6-phosphate dehydrogenase. Thus, the comparison resolves agreement at specific reactions and identifies alternative carbon-routing assumptions for further refinement.

Enzyme-capacity and thermodynamic constraints improved predictions of growth and respiratory physiology against a common experimental dataset from aerobic, glucose-limited chemostats of *A. niger* NW185 (Fig. 2D–G) [21]. All three constrained models reduced errors in growth rate, oxygen uptake, carbon dioxide release and respiratory quotient relative to FBA1015. ecTFA had the lowest errors across all four quantities within our model family, reducing oxygen-uptake and carbon-dioxide-release RMSE by approximately 49% and 50%, respectively. Its respiratory-quotient RMSE (0.068) was the lowest among all eight models, whereas iDU1756 gave the lowest errors for growth and the two gas-exchange rates. GECKO light and ecTFA also approached a finite growth ceiling near 0.18 h*^−^*^1^ as glucose supply increased, reflecting the enzyme-capacity calibration described in Methods 3.2. Together, these results show that the additional constraints improve the joint representation of growth and gas exchange within the shared network.

Citrate production envelopes extend the platform from growth prediction to analysis of growth–production trade-offs (Fig. 2H). At a fixed glucose uptake of 1.83 mmol gDW*^−^*^1^ h*^−^*^1^, maximum citrate secretion declined with increasing growth, and the added constraints narrowed the attainable production range. The black point marks measured citrate secretion by the glucoamylase-producing strain DS03043 in the published batch-culture study [20]. This condition supplies an experimentally measured glucose-uptake rate for the simulations; neither the strain nor the cultivation conditions were selected here for optimized acid production. The envelopes therefore describe computational production capacity under a measured substrate supply, with the black point locating the observed physiological state within that capacity.

### 2.3 Genome-wide coverage and insertion biases of the *Ac/Ds*

The *Ac/Ds* system enabled genome-wide insertion profiling in rich medium and on glucose or xylose as the sole carbon source (Fig. 3A). The analysed libraries are designated DPY1/DPY2 (rich medium), G1/G2 (glucose) and X1/X2 (xylose), with numerals 1 and 2 distinguishing the two libraries under each condition. Induced libraries were propagated under each condition and recovered junctions were mapped to the ATCC 1015 genome (Methods 3.4 and 3.5.1).

**Fig. 3.**
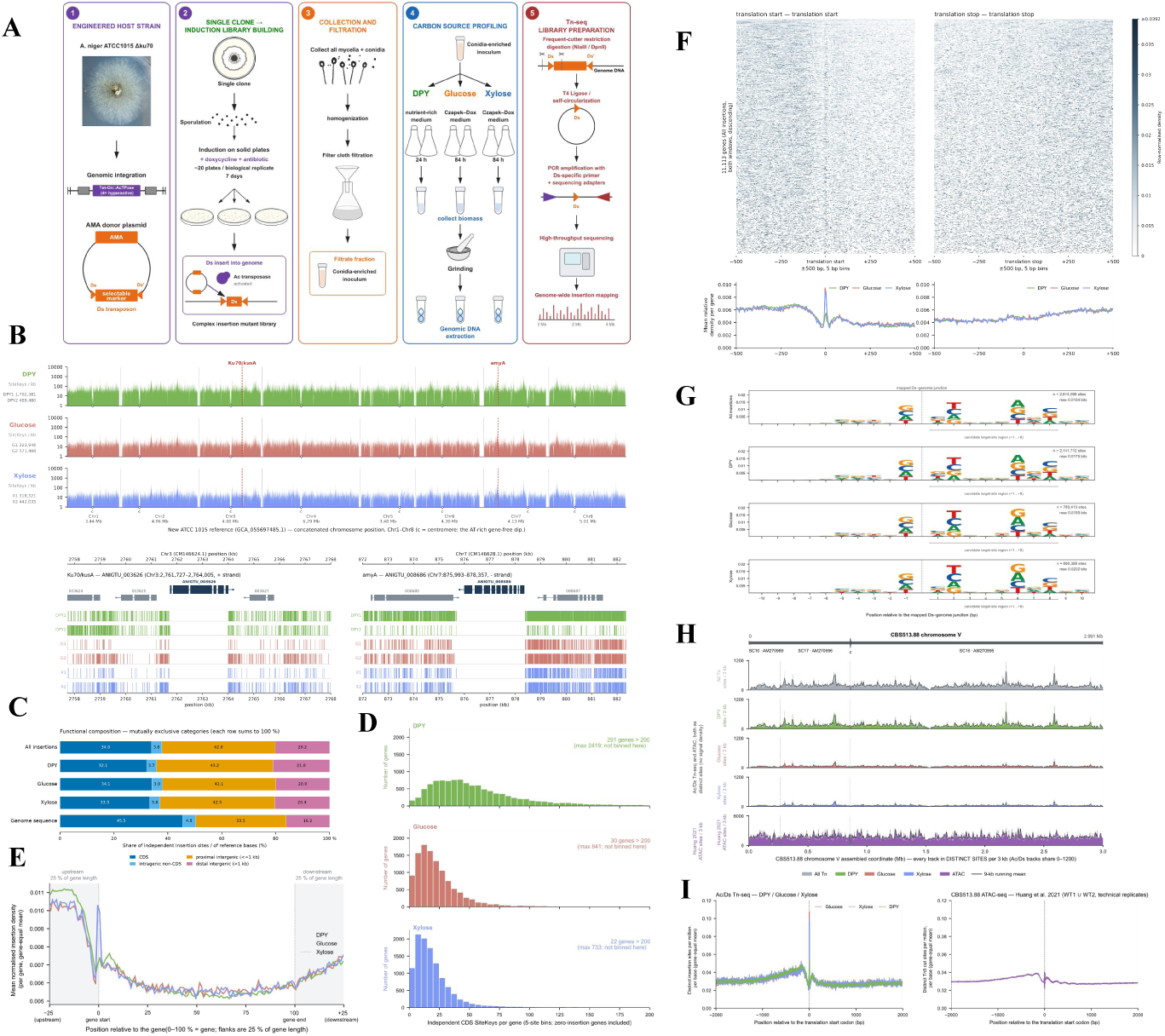
Coverage and insertion biases of the *Ac/Ds* libraries. (**A**) Library construction and growth-condition workflow. DPY denotes rich medium; G and X denote defined media with glucose and xylose, respectively, as the sole carbon source. Numerals distinguish the two analysed libraries per condition. (**B**) Distinct insertion sites per 1 kb window on a log scale. Markers denote candidate centromeric insertion minima; engineered *ku70* and *amyA* loci are shown below. (**C**) Genome and insertion composition across four mutually exclusive classes: coding, intragenic non-coding, proximal intergenic (within 1 kb) and distal intergenic. Pooled distributions combine all six libraries; condition-specific distributions combine their two replicates, deduplicating chromosome–position–strand sites. (**D**) Distinct coding-sequence sites per gene, combining both replicates within each condition. (**E**) Insertion density across the spliced coding sequence and flanks extending 25% of its length. Counts are divided by each gene’s total across this window before averaging with equal gene weights; zero-count genes remain zero. (**F**) Translation-start and -stop profiles in 5 bp bins within *±*500 bp. Each row represents one gene, normalized by its total insertions in the displayed window and ordered by pooled counts across both start and stop windows; zero-count rows remain zero. Heatmaps pool all six libraries; colour is capped at the displayed limit. (**G**) Sequence logos for 20 bp around mapped *Ds*– genome junctions. (**H**) Pooled insertions and published ATAC-seq [19] on CBS 513.88 chromosome V. Insertion counts use 3 kb bins smoothed over 9 kb; tracks have separate vertical scales. The centromere marker denotes the published chromosome-arm junction [22]. (**I**) Translation-start profiles expressed as fractions of each dataset’s site total, with separate vertical scales.

The libraries combined dense genome-wide coverage with localized regions of insertion depletion (Fig. 3B). Across the two replicate libraries analysed for each condition—rich DPY medium and defined media containing glucose or xylose as the sole carbon source—whole-genome densities for individual libraries ranged from one distinct insertion site per 21–112 bp across all eight chromosomes. Each library reached at least 99.1% of annotated gene regions, defined as gene bodies and their 1 kb flanks. Against this broadly sampled background, one AT-rich, gene-poor region on each chromosome showed 9- to 15-fold lower insertion density than the remainder of the corresponding chromosome. These features, together with positional support on six chromosomes from the published CBS 513.88 physical map [22], support their assignment as candidate centromeric regions. Regional composition, insertion depletion and alignment evidence are summarized in Supplementary Table S1.

Broad coding-sequence coverage coexisted with enrichment of insertions in non-coding and gene-terminal regions. Coding sequence comprised 45.5% of the assembly but contained 34.0% of distinct sites pooled across all six libraries, while each non-coding category was proportionally overrepresented (Fig. 3C). Combining the two replicates within each condition, median coding-sequence counts were 47 sites per gene in DPY, 19 on glucose and 15 on xylose; the proportion of genes lacking coding insertions remained below 1.6% in each condition (Fig. 3D). Gene-normalized profiles showed enrichment near both termini and depletion toward gene centres, with similar patterns on glucose and xylose and lower relative density around the 5*^′^*region in DPY (Fig. 3E). Local profiles resolved a sharp translation-start peak on glucose and xylose; DPY also showed start-codon enrichment, although its maximum lay upstream. At the 3*^′^* end, insertion density increased from the coding sequence into the downstream flank (Fig. 3F). Thus, high gene coverage was accompanied by a shared positional bias within and around genes.

The insertion landscape showed little sequence preference but a clear association with chromatin accessibility. Sequence logos across the 20 bp junction window revealed no pronounced insertion motif, with a maximum information content of only 0.016 bits per position (Fig. 3G). To examine whether chromatin organization contributed to the spatial pattern, we compared insertion density with published CBS 513.88 ATAC-seq data [19], using junction reads remapped to CBS 513.88 and counting both signals in the same 3 kb windows. The pooled Spearman correlation was 0.63 (95% block-bootstrap interval, 0.60–0.65), with positive associations on every chromosome (Fig. 3H; Supplementary Table S2). Gene-centred profiles further showed that both signals peaked near translation starts, although ATAC-seq also displayed a broader upstream enrichment (Fig. 3I). The correspondence at both chromosomal and gene-centred scales is consistent with preferential recovery of insertions in accessible chromatin.

### 2.4 Insertion tolerance and essentiality prediction across carbon sources

Insertion-absence screening identified few candidates shared across all conditions (Fig. 4A). After excluding poorly mappable genes and the engineered loci, four genes lacked insertions in the central 80% of their coding sequence across all six libraries, while another 41 lacked them on glucose and xylose but contained them in DPY. These groups required nearby flanking insertions to distinguish gene-level absence from broader unsampled regions (Methods 3.5.2). The four universally insertion-free candidates were short (276–555 bp), limiting the evidence provided by zero counts alone.

**Fig. 4.**
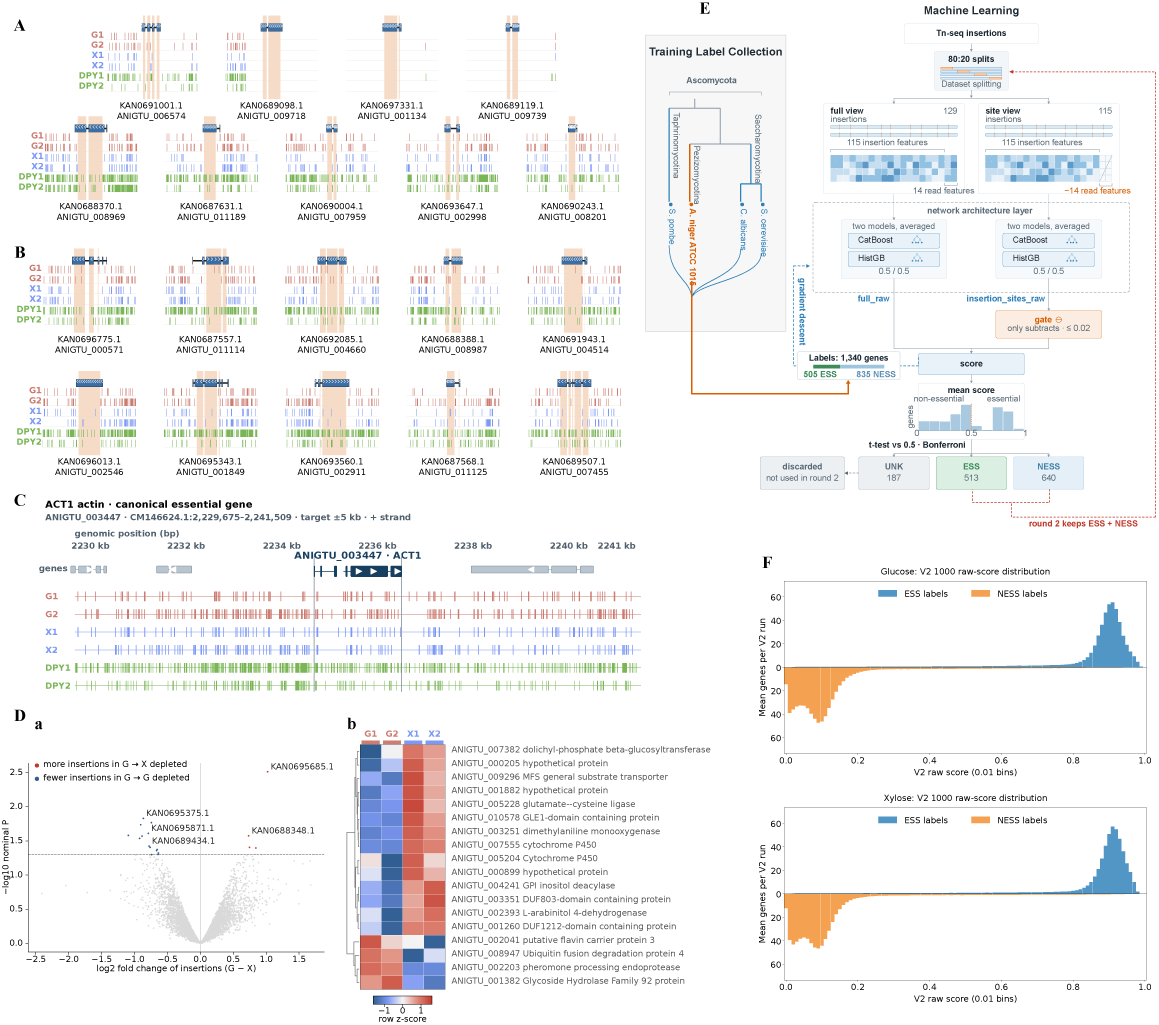
Insertion tolerance and essentiality prediction. (**A**) Insertion tracks for four genes lacking insertions in the central 80% of coding sequence in all six libraries (upper), and five examples from 41 genes lacking them on glucose and xylose but containing them in DPY (lower). Both groups require at least one glucose/xylose insertion in the external 1 kb flanks. Genes with poor sequence mappability and the two engineered loci were excluded (Methods 3.5.2). DPY, rich medium; G/X, glucose/xylose as sole carbon source; numerals identify the two analysed libraries per condition. (**B**) Five displayed examples from each carbon-source-specific group: 76 genes without central coding insertions on xylose but with insertions on glucose (upper), and 38 with the opposite pattern (lower). Presence requires at least one insertion in either replicate; these groups have no flank requirement. Examples in A–B follow the source analysis ranking; peach shading marks the central 80% of coding sequence, with 1 kb flanks shown separately. (**C**) The actin orthologue ANIGTU_003447 and its 5 kb flanks. (**D**) Glucose–xylose differential insertion analysis: (**a**) estimated log_2_(glucose/xylose) fold change in distinct coding-exon insertion-site counts versus *−* log_10_ *P* for 6,289 genes; (**b**) the 18 genes with nominal *P <* 0.05, shown as row *z*-scores of voom log_2_ counts per million. (**E**) Classifier workflow: V1 label selection followed by V2 repeated fitting (Methods 3.5.4). G/X identify glucose/xylose in the V1 score-class counts; V2 retains the original yeast-derived ESS/NESS labels of selected genes, not their V1 score classes. (**F**) Raw-score distributions across 1,000 fits in V2 repeated fitting; higher scores indicate greater support for essentiality. ESS/NESS denote essential/nonessential reference labels transferred from yeast. Bar heights are mean counts per fit in 0.01-wide bins; ESS labels are above zero and NESS labels below. Following V1 label selection, retained sets comprise 504 ESS and 649 NESS labels on glucose, and 504 ESS and 647 NESS labels on xylose. Scores include both training and test predictions, rather than per-gene means; test-only criterion performance is reported in Table 2.

**Table 2.** Agreement of stable essentiality predictions with yeast-derived reference labels using test-only scores.

| Metric | Glucose | Xylose |
| --- | --- | --- |
| Held-out predictions per gene, range | 162–249 | 163–250 |
| Stable ESS / NESS calls, n | 171 / 308 | 201 / 337 |
| Unclassified genes (UNK), n | 674 | 613 |
| Classified genes, n/N (%) | 479/1,153 (41.54) | 538/1,151 (46.74) |
| Accuracy (%) | 91.44 | 90.33 |
| ESS precision (%) | 90.64 | 89.55 |
| ESS recall (%) | 86.11 | 85.31 |
| MCC | 0.8164 | 0.7962 |

ESS/NESS denote essential/nonessential; UNK denotes unclassified. After V1 label selection removed genes with inconclusive score rankings, V2 repeated fitting used 1,000 training/test splits. Each gene was evaluated only in splits in which it was excluded from training. Coverage (n/N) uses all selected reference genes; accuracy, ESS precision, ESS recall and Matthews correlation coefficient (MCC) use only stable ESS/NESS predictions. Evaluation is conditional on prior V1 label selection (Methods 3.5.4).

Carbon-source-specific absence of insertions identified 114 candidate loci (Fig. 4B). Seventy-six lacked central coding-region insertions on xylose but contained them on glucose, and 38 showed the reverse pattern. These patterns nominate carbon-source-associated differences in insertion tolerance, but absence alone does not establish essentiality.

Insertions at the actin locus show why insertion recovery cannot establish dispensability (Fig. 4C). ANIGTU 003447 contained coding insertions in all six libraries despite the essentiality of its yeast orthologue [23]. A plausible explanation is heterokaryotic complementation: intact copies in other nuclei can support the shared cytoplasm, allowing nuclei carrying essential-gene disruptions to persist in *Aspergillus* [18, 24].

Differential insertion analysis likewise provided limited evidence for carbon-source-specific requirements (Fig. 4D). Of 6,289 genes retained after count filtering, 18 showed differences in distinct coding-exon insertion-site counts at nominal *P <* 0.05, but none passed BH-adjusted *P <* 0.05. Together, the sparse insertion-absence candidates, insertion tolerance at a conserved essential locus and limited differential signal motivated a machine-learning approach that combines several features of the insertion landscape. Such an approach can accommodate informative insertion patterns even when heterokaryotic complementation or partial retention of gene function makes absence alone unreliable.

A machine-learning classifier combined insertion patterns, read distributions and gene structure to predict essentiality across 11,111 genes (Fig. 4E). Conserved yeast essentiality supplied reference labels; functional annotations and metabolic-model predictions were excluded from the predictors (Methods 3.5.4). The workflow comprised V1 label selection, which removed reference genes with inconclusive score rankings, and V2 repeated fitting, which used the selected genes with their original reference labels. Across 1,000 V2 fits, essential (ESS) and nonessential (NESS) reference labels concentrated toward opposite ends of the score range, with higher scores indicating greater support for essentiality (Fig. 4F). Stable ESS predictions required a mean score above 0.8 and at least 90% of scores above 0.5; stable NESS predictions required a mean below 0.2 and at least 90% below 0.5. Table 2 evaluates this criterion using only scores obtained when a reference gene was excluded from V2 training, conditional on prior V1 label selection. Genome-wide classification instead used all 1,000 V2 scores per gene, including training-fit scores for reference genes. The criterion selected predictions with high agreement to the reference labels while leaving uncertain genes unclassified, providing a defined set for model comparison.

### 2.5 Comparison of stable essentiality predictions with metabolic models

Genes with stable ESS predictions on both glucose and xylose were enriched for ribosome assembly and RNA maturation (Fig. 5A). Ribosome biogenesis and rRNA processing were among the strongest enrichments, alongside related processes in RNA metabolism and gene expression. These overlapping terms indicate a common concentration of predicted requirements in the cellular machinery for growth.

**Fig. 5.**
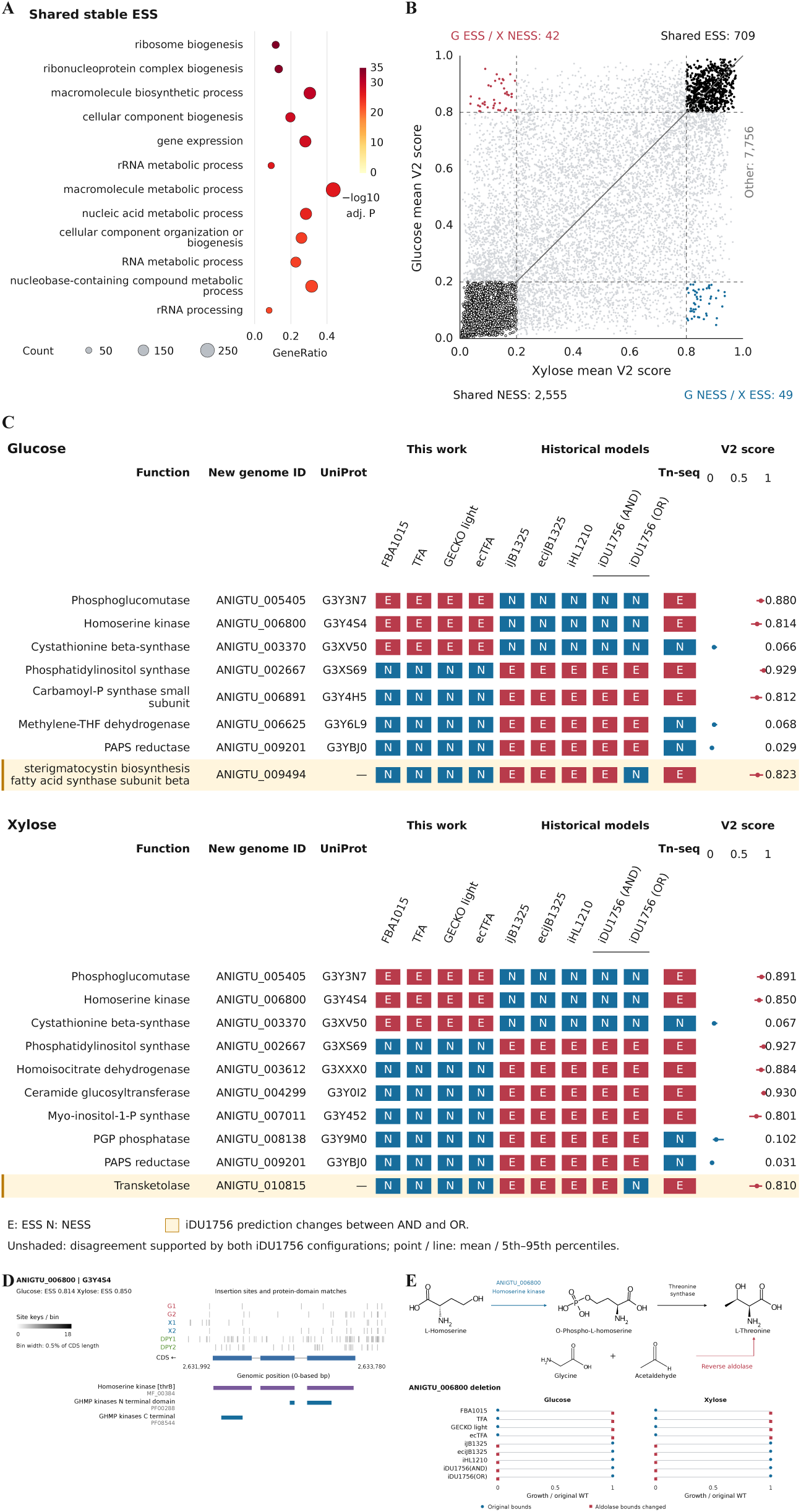
Stable essentiality classifications and their agreement with metabolic models. Detailed legend on the following page. (A) The 12 biological-process terms with the smallest globally BH-adjusted *P* values for shared stable ESS genes. Of 709 genes predicted essential on both glucose and xylose, 603 had GO annotations and form the foreground. GeneRatio is foreground hits/603; point area represents hit count and colour represents *−* log_10_(adjusted *P*). The background universe comprises these 603 genes plus 1,827 other genes with stable predictions in both conditions and GO annotations. Correction covers all 5,158 tests across annotation layers and comparison groups (Methods 3.5.5). (B) Mean classification scores across 1,000 fits in V2 repeated fitting for 11,111 genes (higher scores support essentiality): xylose on the *x*-axis and glucose on the *y*-axis. The diagonal is *y* = *x*; dashed thresholds are 0.2 and 0.8. Solid black, shared stable ESS; hollow black, shared stable NESS; red, G ESS/X NESS; blue, G NESS/X ESS; grey, other genes. ESS/NESS, predicted essential/nonessential. Stable predictions also require at least 90% of scores above/below 0.5, respectively, in addition to the mean-score thresholds. (C) Gene–condition disagreements between the four models developed here and published comparison models. Adjacent iDU1756 (AND) and iDU1756 (OR) columns assign AND or OR only to its unspecified multi-gene associations; explicit gene rules are preserved. The 16 unshaded records require agreement across both configurations; yellow indicates “iDU1756 prediction changes between AND and OR.” E, ESS; N, NESS. Points and lines show means and 5th– 95th percentiles across 1,000 classifier fits, not confidence intervals. A dash indicates no accepted UniProt mapping. (D) ANIGTU 006800 insertion sites, coding exons, the selected HAMAP signature and all Pfam matches on a common genomic axis. Bins span 0.5% of CDS length with a log(1 + count) grayscale; the arrow marks the negative strand. Full domain descriptions are retained. (E) Threonine-pathway scheme and growth after ANIGTU 006800 deletion normalized to each configuration’s original-bound wild-type optimum. Circles show original aldolase bounds; squares show reverse aldolase allowed in the work models or aldolase blocked in the historical models, including both iDU1756 configurations. Original-bound normalization permits ratios above one. Auxiliary substrates and products are omitted; arrows represent the computational mechanism tested.

Stable classifications distinguished shared essentiality predictions from carbon-source-associated candidates (Fig. 5B). Glucose and xylose shared 709 ESS and 2,555 NESS predictions, while 91 genes had opposing classifications (Supplementary Table S3). Only 11 of these 91 genes were mapped to any metabolic model, and none of the models representing them predicted the corresponding reversal.

Model performance was assessed against stable insertion-derived ESS/NESS predictions by simulating gene deletions; a growth rate below 1% of the corresponding wild type defined a model ESS prediction (Methods 3.5.6). Agreement varied with the evaluable gene population (Table 3). Across eight original models represented by nine configurations, eciJB1325 had the highest MCC on each model’s own mapped set under both carbon sources. The two iDU1756 configurations differed only in the AND or OR assignment to unspecified multigene associations and were evaluated on identical genes.

**Table 3.** Agreement of simulated gene-deletion predictions with stable insertion-derived essentiality predictions, using each configuration’s own mapped genes with stable reference predictions and resolved solver outcomes. ESS/NESS counts describe the insertion-derived reference; ESS (essential) is the positive class and NESS denotes nonessential. Model ESS requires deletion growth below 1% of wild type; metrics are point estimates. Nine configurations represent eight original models. The iDU1756 AND/OR assignments concern only unspecified multigene associations, preserving explicit rules. Both rows use identical genes within each carbon source, including genes with different AND/OR predictions. Full-precision metrics and confusion counts are supplied as source data.

| Model | Glucose |  |  |  | Xylose |  |  |  |
| --- | --- | --- | --- | --- | --- | --- | --- | --- |
|  | n (ESS/NESS) | Precision | Recall | MCC | n (ESS/NESS) | Precision | Recall | MCC |
| FBA1015 | 536 (162/374) | 0.758 | 0.154 | 0.254 | 619 (222/397) | 0.767 | 0.149 | 0.233 |
| TFA | 536 (162/374) | 0.735 | 0.154 | 0.245 | 619 (222/397) | 0.750 | 0.149 | 0.226 |
| GECKO light | 536 (162/374) | 0.758 | 0.154 | 0.254 | 619 (222/397) | 0.767 | 0.149 | 0.233 |
| ecTFA | 536 (162/374) | 0.735 | 0.154 | 0.245 | 619 (222/397) | 0.750 | 0.149 | 0.226 |
| iJB1325 | 476 (161/315) | 0.735 | 0.224 | 0.284 | 529 (194/335) | 0.742 | 0.237 | 0.284 |
| eciJB1325 | 462 (152/310) | 0.766 | 0.237 | 0.313 | 525 (190/335) | 0.746 | 0.247 | 0.295 |
| iHL1210 | 409 (141/268) | 0.602 | 0.355 | 0.274 | 468 (174/294) | 0.593 | 0.293 | 0.217 |
| iDU1756(AND) | 612 (187/425) | 0.512 | 0.332 | 0.223 | 692 (235/457) | 0.500 | 0.298 | 0.171 |
| iDU1756(OR) | 612 (187/425) | 0.570 | 0.262 | 0.232 | 692 (235/457) | 0.576 | 0.226 | 0.196 |

On the fixed intersection of genes mapped and evaluable in every configuration, the strongest agreement depended on the carbon source (Table 4). FBA1015 and GECKO light had the highest MCC point estimate on glucose (0.306), whereas iJB1325 ranked highest on xylose (0.312), closely followed by FBA1015 and GECKO light (0.308). The latter two had the highest precision in both conditions. Enzyme constraints left binary predictions unchanged within the FBA1015–GECKO light and TFA–ecTFA pairs.

**Table 4.** Agreement of simulated gene-deletion predictions with stable insertion-derived essentiality predictions on fixed common gene sets: 243 glucose genes (88 ESS, 155 NESS) and 270 xylose genes (110 ESS, 160 NESS), mapped and evaluable in all nine configurations. Counts describe the insertion-derived reference. ESS/NESS denote essential/nonessential; model ESS requires deletion growth below 1% of wild type. The iDU1756 AND/OR assignments concern only unspecified multigene associations, preserving explicit rules; genes with different predictions are retained. ESS is the positive class; metrics are point estimates. Full-precision metrics, confusion counts and memberships are supplied as source data.

| Model | Glucose |  |  | Xylose |  |  |
| --- | --- | --- | --- | --- | --- | --- |
|  | Precision | Recall | MCC | Precision | Recall | MCC |
| FBA1015 | 0.778 | 0.239 | 0.306 | 0.800 | 0.255 | 0.308 |
| TFA | 0.750 | 0.239 | 0.291 | 0.778 | 0.255 | 0.296 |
| GECKO light | 0.778 | 0.239 | 0.306 | 0.800 | 0.255 | 0.308 |
| ecTFA | 0.750 | 0.239 | 0.291 | 0.778 | 0.255 | 0.296 |
| iJB1325 | 0.733 | 0.250 | 0.290 | 0.775 | 0.282 | 0.312 |
| eciJB1325 | 0.733 | 0.250 | 0.290 | 0.756 | 0.282 | 0.300 |
| iHL1210 | 0.538 | 0.318 | 0.191 | 0.564 | 0.282 | 0.161 |
| iDU1756(AND) | 0.531 | 0.386 | 0.210 | 0.554 | 0.373 | 0.183 |
| iDU1756(OR) | 0.533 | 0.273 | 0.170 | 0.566 | 0.273 | 0.160 |

Gene-level comparisons exposed opposing predictions of gene requirements (Fig. 5C). Sixteen gene–condition records across 11 genes showed opposing predictions between the four models developed here and the four published comparison models, requiring agreement under both iDU1756 configurations. The stable insertion-derived predictions supported each model group in eight records. Two additional records, highlighted in yellow, changed prediction between iDU1756 AND and OR and were excluded from this unanimous-disagreement count.

ANIGTU 006800 linked a consistent model disagreement to a plausible functional reassignment (Fig. 5D). It was selected because stable ESS predictions supported all four work models on both carbon sources, whereas every historical configuration predicted NESS. Although annotated as a trihydroxynaphthalene reductase, the protein carried a near-full-length HAMAP homoserine-kinase signature and a strong N-terminal GHMP-kinase match; the C-terminal Pfam match was weaker. These signatures support the homoserine-kinase function represented in the metabolic models, linking this locus to the conventional threonine-biosynthesis pathway. The essentiality disagreement therefore prompted examination of routes bypassing that pathway. Coding insertions occurred in all six libraries, illustrating that its stable ESS classification did not require an insertion-free gene.

An alternative route to threonine explained the opposing deletion phenotypes computationally (Fig. 5E). Reverse threonine-aldolase flux can supply threonine from glycine and acetaldehyde, bypassing homoserine kinase in the models. This direction was available in the historical models but blocked by the original bounds of the four models developed here. Blocking aldolase abolished ANIGTU 006800-deletion growth in every historical configuration, while allowing reverse aldolase flux restored growth to approximately the original wild-type level in all four work models. This intervention identifies a pathway assumption for testing, without establishing bypass activity in vivo.

## 3 Methods

### 3.1 Reconstruction and curation of FBA1015

FBA1015 was reconstructed from the ATCC 1015 proteome using the RAVEN homology-based protocol [25]. The templates were yeast-GEM for *Saccharomyces cerevisiae* [26] and rhto-GEM for *Rhodotorula toruloides* [27]. The network combined homology-supported reactions, gap-filling hypotheses and curated additions. The published *A. niger* model iJB1325 supplied biomass composition and additional network content [4]; subcellular localisation was predicted with DeepLoc 2.1 [28].

#### 3.1.1 Homology-based reconstruction and gap filling

Reciprocal proteome searches used an initial E-value threshold of 10*^−^*^4^. Hits were passed to RAVEN’s *getModelFromHomology* [29] with an E-value threshold of 10*^−^*^30^, minimum alignment length of 150 amino acids and minimum identity of 35 %. Template gene rules were evaluated after removal of unmatched genes; reactions were retained when a complete alternative supported by target genes remained. Transferred rules contained only ATCC 1015 identifiers. Gap-filled reactions received gene rules only when supported by curated target-specific records.

Gap filling first used MENECO [30] with the *S. cerevisiae* template, followed by RAVEN, minimizing the number of added donor reactions. Biomass precursors were evaluated individually, followed by the complete biomass reaction if simultaneous precursor production remained insufficient. The *S. cerevisiae* template was used first and the *R. toruloides* template for unresolved requirements. Additions were accepted only if the resulting network produced the target. This procedure was repeated for D-glucose, D-xylose, L-arabinose, D-arabinose, D-mannose, maltose and glycerol. Target production refers to the tested precursor drain or complete biomass reaction; the source package supplies the stage-specific production thresholds and medium bounds (Supplementary Methods D).

#### 3.1.2 Biomass composition and lipid representation

Biomass equations followed iJB1325 [4]. Lipid metabolism was rebuilt using SLIMEr [31], which represents lipid classes and acyl-chain composition through pooled components. Coefficients were derived from measured *A. niger* V35 lipid abundances [32], used as a surrogate for ATCC 1015.

#### 3.1.3 Network integration with iJB1325

Before integration, iJB1325 locus tags were mapped to ATCC 1015 genes and its compartments were expanded using DeepLoc predictions of protein localization and membrane association. For a non-transport reaction with gene associations, predicted compartments of its associated proteins defined candidate reaction locations; reactions without localization evidence retained their original placement. The compartment definitions and curated exceptions are documented in Supplementary Methods D. Metabolites were merged only when supported by a shared structural identifier or an exact name with matching formula, charge and compartment; formula alone was insufficient. The merged network was manually curated. A separate reconstruction based on KEGG orthology contributed 35 selected reactions. Reaction provenance records distinguish template-derived, iJB1325-derived, KEGG-derived and manually constructed interface reactions.

### 3.2 Construction of enzyme- and thermodynamically constrained models

TFA, GECKO light and ecTFA were derived from the fixed FBA1015 network by adding thermodynamic constraints, enzyme costs, or both. All four models share the same biochemical network and gene set, allowing each constraint layer to be compared with and without the other.

#### 3.2.1 Shared network and parameterization

FBA1015 defined the shared biochemical network for all models. Temperature was 298.15 K and total protein content was 0.263 g gDW*^−^*^1^. The enzyme mass fraction, *f* = 0.4189, was estimated from measured *A. niger* glucose-grown proteomic abundances [33] as the mass fraction represented by model enzymes. Enzyme saturation, *σ*, was fitted with the GECKO routine to 0.18 h*^−^*^1^, the upper end of the reported ATCC 1015 maximum growth rate of 0.17 *±* 0.01 h*^−^*^1^ [34]. Glucose supply was relaxed during fitting because the assay-medium bound prevented the target growth rate. Fits at glucose supplies of 1.5, 2 and 3 mmol gDW*^−^*^1^ h*^−^*^1^ returned *σ* = 0.92, 0.87 and 0.87, respectively; *σ* = 0.87 was used throughout. The resulting enzyme pool, *P*_tot_*fσ*, was 95.85 mg gDW*^−^*^1^.

#### 3.2.2 Enzyme constraints in GECKO light

GECKO 3 light [8] constrained total enzyme usage through a shared protein pool, without imposing individual enzyme-abundance bounds. Gene rules were converted to disjunctive normal form to enumerate isozyme and complex candidates. For each reaction direction, enzyme cost was calculated as the sum of subunit molecular weights weighted by copy number, divided by turnover number and 3,600 s h*^−^*^1^. The least costly eligible candidate supplied the pool coefficient.

Turnover numbers were assigned in the following order: substrate-specific BRENDA values, DLKcat predictions from substrate structure and enzyme sequence [35], lower-specificity BRENDA values, and the GECKO default. Candidate-level assignments and their sources were retained in the enzyme-parameter tables.

#### 3.2.3 Thermodynamic constraints in TFA

Thermodynamic constraints were implemented with pyTFA [7]. Reference pH, ionic strength, metabolite-concentration bounds and relative electrical potentials were assigned to eight aqueous compartments; six non-aqueous compartments received no aqueous thermodynamic constraints. Compartment identities and parameter bounds are supplied in Supplementary Methods D. Gibbs-energy constraints were available for 1,187 reactions. Directional feasibility was evaluated over the bounds of formation-energy and concentration variables, including their uncertainty, rather than from nominal standard Gibbs energies alone. To restore growth to at least 1 % of the FBA1015 optimum in the defined glucose reference medium, the lower bound of the standard Gibbs energy for argininosuccinate lyase was relaxed by the minimum required amount, 2.46 kcal mol*^−^*^1^. Other conflicts were recorded without adjustment.

#### 3.2.4 Combined enzyme and thermodynamic constraints in ecTFA

The enzyme-pool constraint was applied directly to the forward and reverse flux variables of the thermodynamic model. Coefficients were checked against GECKO light before and after gene deletion. Deletions removed every enzyme candidate containing the deleted gene, and costs were recalculated from the least costly surviving isozyme; directions without a surviving candidate were closed.

### 3.3 Model evaluation against published datasets

Models were evaluated using published phenotype, intracellular-flux and physiological datasets. These benchmarks were not used to fit the four models developed here. However, they had been used in developing or evaluating the historical models, and iJB1325 contributed reconstruction evidence to FBA1015; the comparisons therefore constitute shared benchmarks rather than independent validation.

The phenotype benchmark comprised the 471 cases encoded in the gem:listOfTests block of iJB1325. Cases were evaluated independently with their specified bounds, gene deletions and expected outcomes; unspecified nutrient uptake was closed. Cases 141–143 required biomass flux below *−*999 despite a nonnegative biomass bound and were excluded from every model, leaving 468 eligible cases. The remaining tests include growth, biomass-component synthesis and specified reaction-flux outcomes, evaluated against their encoded thresholds with a 10*^−^*^8^ tolerance. In cases 126, 128 and 417, a negative-flux expectation applied to a reaction with a zero lower bound was interpreted as no growth. Cross-model metabolite matching required a shared structural identifier or normalized name, together with matching heavy-atom composition and compartment. Biomass drains were added to iDU1756 and iHL1210 on their native biomass metabolites; single-metabolite interfaces were also added for K^+^, Ca^2+^ and sulfate, which lacked uptake reactions. The iHL1210 gene-deletion cases remained unresolved because the benchmark mapping did not translate its AspGD identifiers to the JGI locus identifiers used by the iJB1325 test definitions. Coverage was the number of resolved cases out of 468; accuracy and Matthews correlation coefficient (MCC) used only those resolved cases. Proven infeasibility was scored as no growth; unresolved cases were excluded.

Intracellular-flux comparisons used two published batch-culture states [20]. Measured growth, glucose uptake, O_2_ uptake, CO_2_ release, citrate secretion and oxalate secretion were fixed as equalities, and unspecified carbon uptake was closed. Parsimonious FBA minimized total absolute biochemical flux [36]; isotope-derived internal fluxes were reserved for evaluation. Twenty-eight carbon transformations were mapped by chemical identity, with the externally constrained oxalate transformation excluded, leaving 27 scored transformations per strain. Fluxes were expressed per 100 mmol glucose and measured values were regressed against predictions. Because the measurements were from CBS 513.88 and DS03043, these comparisons assess conditional internal-flux agreement across strains.

Growth was maximized at each glucose-uptake ceiling, after which pFBA selected gas-exchange fluxes while retaining growth within 10*^−^*^8^ of the optimum. The resulting curves were compared with four published observations from aerobic, glucose-limited chemostats of *A. niger* NW185 [21]. Citrate production envelopes were calculated separately by maximizing citrate secretion [6] at prescribed growth rates and a fixed glucose uptake of 1.83 mmol gDW*^−^*^1^ h*^−^*^1^, matching the DS03043 condition of [20]. The corresponding measured growth and citrate-secretion rates were shown as a reference point. This glucoamylase-production dataset provided an experimentally observed glucose uptake, rather than a condition optimized for citrate production.

### 3.4 Construction of the *Ac/Ds* insertion library

#### 3.4.1 Host strain and transposon constructs

The recipient was *Aspergillus niger* ATCC 1015 with *ku70* (ANIGTU 003626) deleted to facilitate targeted integration. AcTPase4x, a hyperactive maize *Ac* transposase carrying four amino-acid substitutions [37], was integrated at *amyA* (ANIGTU 008686). A Tet-on cassette [38, 39] controlled expression and comprised the *A. nidulans gpdA* promoter, rtTA2S-M2, the *cgrA* terminator and seven *tetO* operators fused to a minimal *gpdA* promoter.

The donor plasmid contained the extrachromosomal AMA1 replication element [40] and a selectable *nat* cassette between the two *Ds* ends. It was introduced by PEG-mediated protoplast transformation.

#### 3.4.2 Transposition induction and library amplification

Transformants were selected on potato dextrose agar (PDA) containing zhongshengmycin, a streptothricin-class antibiotic inactivated by the *nat* product [41]. A mycelial rim with its underlying agar was transferred to fresh selective PDA and incubated for seven days to produce conidia.

Conidia were spread at approximately 7 × 10^6^ per plate onto 20 PDA plates containing 2,500 *µ*g mL*^−^*^1^ zhongshengmycin and 20 *µ*g mL*^−^*^1^ doxycycline [38]. Each plate was overlaid with a 10-kDa semi-permeable membrane and incubated in darkness for seven days. Biomass was collected from the membranes, dispersed with a handheld homogeniser, filtered through Miracloth (Calbiochem; 22–25 *µ*m), washed and stored in 30% glycerol. Each 20-plate batch constituted one library induction batch.

#### 3.4.3 Culture conditions and sampling

For each culture replicate, 1,000 mL medium was distributed among ten flasks to maintain an inoculum of approximately 10^6^ conidia mL*^−^*^1^. Mycelia from the ten flasks were pooled before DNA extraction. Media were DPY (2% glucose, 1% peptone, 0.5% yeast extract, 0.5% KH_2_PO_4_ and 0.05% MgSO_4_*·*7H_2_O; also termed YPD) [19], and Czapek–Dox (CD) medium with its sucrose replaced by 20 g L*^−^*^1^ glucose or xylose. All media contained 2,500 *µ*g mL*^−^*^1^ zhongshengmycin and no doxycycline.

DPY cultures were harvested at 24 h and CD cultures at 84 h. The longer defined-medium incubation accommodated slower growth and was intended to allow depletion of insertion-bearing nuclei otherwise maintained by heterokaryotic complementation [18].

#### 3.4.4 DNA extraction and junction-sequencing library preparation

Mycelium was vacuum-filtered, freeze-dried at *−*40 °C for 48 h and ground in liquid nitrogen. Genomic DNA was extracted with a commercial kit, treated with 250 *µ*g mL*^−^*^1^ RNase A, re-extracted and quantified by agarose-gel densitometry against a 1 kb ladder using ImageJ.

Junction libraries followed the SATAY protocol [42]. Separate 2 *µ*g DNA aliquots were digested with 50 U NlaIII or DpnII in 50 *µ*L for 16 h at 37 °C, followed by heat inactivation at 65 °C for 20 min. Each digest was processed separately: 48 *µ*L was circularised with T4 DNA ligase in 400 *µ*L, purified, and amplified with transposon-junction primers carrying sequencing adaptors and sample indices. The two preparations were pooled equally for sequencing. Libraries were prepared in-house and sequenced by BGI Genomics.

### 3.5 Insertion-data analysis and essentiality prediction

#### 3.5.1 Read processing and insertion-site mapping

A custom pipeline (Supplementary File 1) retained FASTQ provenance by batch and condition. Reads required an exact *Ds* end tag (5*^′^*-TTTTACCGACCGTTACCGACCGTTTTCATCCCTA-3*^′^*) within the first 100 nt. The genomic sequence following the detected Ds tag was trimmed at the 3*^′^* end below Phred 15 and retained if at least 25 nt long. Bowtie2 alignment used end-to-end, very-sensitive mode. Alignments were excluded for mapping quality *<* 20, tied best positions, or clipping or indels at the insertion junction.

Sites were keyed by chromosome, 1-based position and strand. Identical coordinates were collapsed, and NlaIII and DpnII sites were merged while retaining digest-specific read counts. No UMI or neighbouring-coordinate correction was used; counts therefore represent distinct junction positions rather than independent transposition events. Sequence mappability was assessed using uniqueness of genomic 25-mers on both strands. The insertion-absence and differential-count screens excluded genes for which fewer than 95% of coding bases were callable by this rule (Supplementary Methods D); mappability was distinguished from whether an insertion was observed. The reference was the chromosome-scale ATCC 1015 assembly GCA 055697485.1, comprising eight chromosomes, 35.6 Mb and 11,113 protein-coding genes [43].

#### 3.5.2 Screening for insertion-free coding regions

Insertion-absence screening used the central 80% of the spliced coding sequence, excluding 10% at each end. Absence in a condition required zero sites in both replicates; presence required at least one site in either replicate. Four sets were defined: absence in all three media; absence on glucose and xylose but presence in DPY; absence on glucose but presence on xylose; and the reverse. The first two sets additionally required at least one glucose/xylose insertion within the external 1 kb flanks, providing evidence that the surrounding region had been sampled. The carbon-source-specific sets had no flank requirement and did not depend on DPY status. Genes with a callable coding fraction below 95% and the two engineered loci were excluded.

#### 3.5.3 Differential insertion analysis between carbon sources

Differential analysis used distinct insertion-site counts within coding exons rather than read counts, reducing the influence of highly represented junctions. Counts were TMM-normalized and analysed with voom and limma empirical Bayes. The design was *∼* 0 + group for six samples across three media, with contrasts of DPY versus glucose, DPY versus xylose, and glucose versus xylose. Non-callable genes were excluded, and edgeR’s filterByExpr retained 6,289 genes. Glucose versus xylose was the primary contrast, with positive log_2_ fold changes indicating higher counts on glucose; nominal and BH-adjusted *P <* 0.05 thresholds were reported separately. DPY comparisons were confounded with experimental batch.

#### 3.5.4 Supervised essentiality prediction from insertion patterns

Essentiality prediction followed the two-pass approach of Billmyre et al. [17]. Predictors described insertion positions, read distributions and gene structure; functional annotations and metabolic-model predictions were excluded. Orthology was used only to transfer reference labels. Labels required concordant published essentiality calls in *Candida albicans*, *Saccharomyces cerevisiae* and *Schizosaccharomyces pombe* [16], together with one-to-one orthology to ATCC 1015 (degree one on both sides). The resulting set contained 1,340 reference genes, including 505 essential (ESS) and 835 nonessential (NESS) labels; engineered loci were excluded. The transferred labels and source-to-target identifiers are supplied in the reference-label table (Supplementary Methods D).

Separate classifiers were fitted for glucose and xylose. Each gene was represented by one vector containing features from both replicates as separate columns, with shared sequence features included once. The full panel comprised 129 features describing insertion number, density, insertion-free gaps, local insertion frequency, the 100 kb neighbourhood, promoters, exon–intron structure, restriction-enzyme effects and read concentration. A reduced panel excluded the 14 read-concentration features. Each panel combined CatBoost and histogram gradient boosting with equal weights. Features were neither scaled nor selected. Four-fold inner cross-validation assigned continuous training weights from one to three based on the absolute difference between each reference label and its held-out inner-fold score; weights rose linearly from one at the median error to three at the 90th percentile, with values outside this range clipped; both learners also used class balancing. Each panel score was the mean ESS-class probability from its two learners, on a 0–1 scale. Higher scores indicate greater support for essentiality. Denoting the full-panel score by *s*_full_ and the reduced-panel score by *s*_site_, the combined classification score was

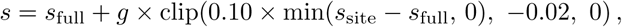

where *g* = 1 for genes meeting the predefined site-support and read-distribution criteria in Supplementary Methods D, and zero otherwise. Thus, the reduced panel could reduce the full-panel score by at most 0.02. The full panel and gate both used read-distribution information; the reduced panel omitted read-concentration features while retaining insertion-position and gene-structure features.

Training used repeated stratified 80/20 splits, with 100 fits for V1 label selection and 1,000 fits for V2 repeated fitting. In V1, predictions for all 1,340 reference genes, including both training and test predictions, were converted to within-fit percentile ranks. A two-sided one-sample *t*-test compared each gene’s mean rank with 0.5, with Bonferroni correction across all labels. Genes with Bonferroni-adjusted *P <* 0.05 and a mean rank above or below 0.5 were retained; other genes were excluded. Retention did not require agreement between this score direction and the original reference label. V1 label selection retained 1,153 genes for glucose and 1,151 for xylose. V2 retained their original transferred labels and generated raw, unranked scores for all 11,111 genes at every fit. Feature definitions, ensemble structure, gate and hyperparameters were fixed. V2 split and fit seeds were 20260911 + 500000 + *i* and 20260911 + 600000 + *i*, respectively, for *i* = 0, … , 999.

The stable-call criterion was evaluated using only predictions from fits in which each retained label belonged to the test set. Each split contained 231 test labels (101 ESS, 130 NESS). Stable ESS required a mean test score *>* 0.8 with at least 90% of test scores *>* 0.5; stable NESS required a mean *<* 0.2 with at least 90% *<* 0.5. Remaining genes were unclassified (UNK). Classification metrics were calculated only for ESS or NESS calls, while coverage used all retained labels as its denominator. Confusion counts, UNK assignments and per-gene test exposure were retained in the source data. Average precision was calculated for each test set and then averaged across fits. This evaluation measures agreement with transferred reference labels after V1 label selection; it is not an independent evaluation of the complete two-stage procedure, because V1 label selection was not repeated within an outer test split.

#### 3.5.5 Stable classification criteria and GO enrichment analysis

Genome-wide stable calls applied the same mean-score and 90% consistency criteria to all 1,000 V2 scores per gene, including training-fit predictions for labelled genes. Scores equal to 0.5 counted toward neither side. UNK genes were excluded from stable-label comparisons. Neither the thresholds nor the classifier architecture was adjusted using model-comparison outcomes.

GO annotations [44] combined InterProScan terms with terms transferred from CBS 513.88 through unique reciprocal best hits requiring *≥* 95% identity and *≥* 90% coverage of both proteins. Terms were propagated through is a and part of relationships in the archived ontology; alternative identifiers were resolved and obsolete or unresolved terms excluded. Shared-ESS enrichment compared 603 annotated shared ESS genes with 1,827 other annotated genes stable in both conditions, giving a universe of 2,430 genes. Eligible terms annotated at least five genes and at most 85% of the annotated foreground-plus-comparison population for that test and annotation layer; ontology roots were excluded. Shared-ESS comparisons used one-sided Fisher exact tests; the other comparisons used two-sided tests.

Benjamini–Hochberg correction was applied jointly to all 5,158 tests in the pre-defined family, covering InterPro-only and combined annotation layers, all three GO namespaces and six comparisons per layer. Comparisons were shared ESS versus other jointly stable genes, the same comparison after excluding all 1,340 original reference-label genes, genes predicted ESS only on glucose versus only on xylose (the other condition could be NESS or UNK), the two opposing ESS/NESS groups, and, separately for each carbon source, stable ESS genes predicted NESS by every model configuration versus the remaining stable ESS genes in the common evaluable set (Methods 3.5.6). The ancillary consensus tests did not determine performance-table membership. Figure 5A displays the 12 combined-layer biological-process terms with the smallest globally adjusted *P* values. Full memberships, contingency counts and the correction family are supplied as source tables.

#### 3.5.6 Gene mapping and model-based essentiality evaluation

Eight original models were evaluated as nine configurations. The four models developed here (FBA1015, TFA, GECKO light and ecTFA) use JGI protein identifiers; iJB1325 JGI transcript identifiers were translated through a typed transcript–protein manifest. iHL1210 and eciJB1325 use CBS 513.88 identifiers, including KEGG-style identifiers where applicable, whereas iDU1756 contains Aspni7 protein and legacy ASPNIDRAFT identifiers. Sequence mapping required *≥* 95% identity, *≥* 90% query and subject coverage, E-value *≤* 10*^−^*^20^ and a unique best target locus. Existing aliases had to agree with sequence evidence; absent aliases and ambiguous split aliases required unique reciprocal best hits within the source sequence namespace. All model genes mapping to one new-genome locus were deleted together. Accepted and unresolved mappings were recorded separately.

Some multigene associations in the published iDU1756 model did not explicitly specify a Boolean relationship. We retained all explicitly defined associations and constructed iDU1756(AND) and iDU1756(OR) by assigning AND or OR, respectively, to each unspecified association.

The published spreadsheet [5] contained 289 unspecified multigene associations, 611 explicit OR associations and 54 explicit AND associations; no convention for interpreting unspecified relationships was identified. Both configurations preserved explicit expression structure, single-gene associations and reactions without genes. They used identical stoichiometry, bounds, objective, medium, mapping and essentiality thresholds. Spreadsheet stoichiometry and bounds were combined with the SBML gene-association tree and biomass objective, retaining the same biomass-demand and sulfate-exchange interfaces in both configurations. iHL1210 maintenance bounds were restored from its published spreadsheet.

Models used nitrate as the nitrogen source and glucose or xylose as the sole carbon source. Unspecified uptake was closed, assay inorganic exchanges were opened, and carbon uptake was capped at 1 for glucose or 1.2 for xylose in mmol gDW*^−^*^1^ h*^−^*^1^, providing equal carbon-atom supply. Exact exchange identities and bounds for each configuration are supplied in Supplementary Methods D. Biomass was maximized for each model and medium. A locus was predicted ESS when *µ*_KO_*/µ*_WT_ *<* 0.01, using that configuration’s positive wild-type optimum under the original assay bounds. Proven infeasibility implied zero growth; missing mappings, unstable reference labels and unresolved solver outcomes were excluded from classification.

Deletions were applied through native gene–protein–reaction rules. GECKO light and ecTFA additionally removed candidates containing the deleted gene, closed directions without surviving candidates and recalculated pool coefficients from the least costly remaining candidate. Native coefficients were checked against the candidate tables. eciJB1325 retained separate isozyme branches and protein constraints; TFA and ecTFA retained native thermodynamic variables and constraints. Gurobi solves used a 30-s limit, a 120-s retry for unresolved outcomes, MIP gap 10*^−^*^7^, and feasibility, optimality and integrality tolerances of 10*^−^*^9^. Identical deletion-effect signatures could share a solve. Wild-type optima were reused only when their primal solutions remained feasible after deletion and coefficient updates; other cases were reoptimized. Model restoration was checked against the wild-type optimum after each deletion.

Table 3 uses each configuration’s mapped, stable-label, solver-resolved population. Table 4 uses their intersection across all nine configurations: 243 genes on glucose (88 ESS, 155 NESS) and 270 on xylose (110 ESS, 160 NESS). The two iDU1756 configurations used identical genes, including those whose predictions differed. ESS was the positive class; precision, recall and MCC were derived from the confusion counts retained in the source tables. Rankings describe point estimates. Figure 5C used a separate selection criterion: the four work models had to agree and oppose iJB1325, eciJB1325 and iHL1210, together with at least one iDU1756 configuration. Agreement across all historical models required both iDU1756 configurations to agree; yellow shading marks a change between AND and OR.

#### 3.5.7 Functional evidence and threonine-aldolase interventions

ANIGTU 006800 insertion and domain tracks used zero-based, half-open genomic coordinates. Distinct chromosome–position–strand sites were projected alongside coding exons, using bins of 0.5% coding-sequence length, flanks of 25% of the spliced coding-sequence length on each side and a log(1 + count) grayscale referenced to 18 sites per bin. InterProScan domain coordinates were projected through the spliced coding sequence, accounting for the negative strand. The panel includes the selected HAMAP family match and all Pfam matches with full descriptions; complete hits are supplied as source data.

The locus deletion was tested under original and altered threonine-aldolase bounds in all nine configurations and both carbon sources, giving 36 simulations. In the four work models, the lower bound of cytosolic threonine aldolase r 1040 was changed from 0 to *−*1000 to allow reverse flux. All native r301 branches in iJB1325 and eciJB1325, and R515 in iHL1210 and both iDU1756 configurations, were closed in both directions. Other constraints were retained, including enzyme-candidate updates and thermodynamic constraints. Growth was normalized to each configuration’s wild-type optimum in the same medium under its original bounds, without truncating ratios at one. These interventions test dependence on reaction bounds rather than establish in vivo flux or biochemical function.

## 4 Discussion

This study connects a curated metabolic network with high-density insertion profiling to investigate gene requirements in *A. niger*. The platform supports comparisons at two levels: matched models reveal the effects of additional constraints, while insertion-derived classifications identify genes for which the models make conflicting predictions. Its practical contribution is the ability to move from genome-wide patterns to specific assumptions about gene assignment and pathway activity.

FBA1015 extends the metabolic and compartmental representation of ATCC 1015 from the community reconstruction of iJB1325 [4]. Deriving four models from one network makes their differences interpretable, although greater network size alone does not establish greater accuracy. Added enzyme and thermodynamic constraints improved agreement with measured growth and gas exchange, consistent with the value of enzyme constraints reported previously [6]. Their benefits were task-dependent: intracellular-flux agreement did not improve consistently, and common-set essentiality MCC point estimates favoured FBA1015 and GECKO light on glucose but iJB1325 on xylose. The growth plateau partly reflects calibration of enzyme saturation to a published maximum growth rate, while the intracellular-flux test fixes measured exchange rates and uses other strains. These benchmarks therefore assess complementary aspects of performance rather than a single hierarchy of model quality.

The insertion libraries provide dense sampling across the genome and within individual genes. Their nonuniform distribution is informative: enrichment near translation starts and association with published ATAC-seq signal from a different strain and experimental setting [19] indicate that local insertion opportunity should be considered when interpreting depletion. The AT-rich, gene-poor insertion minima also support candidate centromeric regions, with positional evidence from the published physical map on six chromosomes. These observations extend the resource beyond essentiality prediction while retaining the distinction between inferred genomic features and experimentally established functions.

Coding insertions at the actin locus illustrate why insertion recovery cannot be equated with gene dispensability. Heterokaryotic complementation can preserve nuclei carrying essential-gene disruptions [18], and position-dependent retention of function offers another explanation [15]. The present data do not resolve these mechanisms or assign a growth rate to each insertion-bearing lineage. They instead motivate an analysis that combines insertion position, local coverage and gene structure rather than relying on zero-insertion rules.

The classifier implements this broader interpretation of insertion evidence, building on supervised approaches developed in other fungi [16, 17]. The consensus criterion achieved approximately 90–91% accuracy among the 42–47% of retained labels it classified using held-out V2 scores, providing a defined subset for subsequent comparison. These test-only scores evaluate genes already selected in V1 label selection, and the genome-wide classifications additionally include training-run scores for labelled genes. Conserved yeast labels are also proxies for requirements in *A. niger*. Thus, score consistency supports prioritization, while independent characterization of *A. niger* genes remains necessary to establish predictive accuracy in the target organism.

The shared and opposing stable classifications nominate different biological questions. Enrichment of shared ESS predictions for ribosome biogenesis and RNA processing is consistent with common growth requirements, whereas opposing classifications identify candidates for carbon-source-dependent requirements. Only 11 of the 91 opposing-classification genes were represented in any model, exposing the limited reach of metabolic reconstructions into the wider genetic response. Conversely, low model recall against the stable reference directs attention to permissive bypasses, incomplete gene rules and missing requirements, alongside possible reference-label errors. Fixing the comparison populations and evaluating both interpretations of the unspecified iDU1756 associations makes these disagreements traceable without treating the classifier as experimental ground truth.

ANIGTU 006800 provides a concrete example of this process. Protein-family evidence supported homoserine kinase despite a different genome annotation, and its predicted deletion phenotype depended on threonine-aldolase availability. Allowing reverse aldolase flux rescued deletion growth in the four new models, whereas closing aldolase abolished deletion growth in the historical models. These computational interventions identify a pathway assumption for biochemical and genetic testing; they do not establish bypass activity in vivo. Resolving such disagreements with targeted experiments would improve both the functional interpretation of insertion profiles and the mechanistic content of the models.

## 5 Conclusion

We established an integrated metabolic modelling and high-density *Ac/Ds* insertion platform for *A. niger* ATCC 1015. Matched models support assessment of added constraints, while machine-learning analysis of insertion patterns prioritizes shared and carbon-source-associated essentiality predictions. Their comparison connects genome-wide screening to specific gene assignments and pathway assumptions. The resulting models, insertion maps and stable classifications provide a resource for targeted functional validation, model refinement and rational strain engineering.

## Appendix A Supplementary Table S1: Candidate centromeric regions

**Table S1 Supplementary Table S1.** Evidence for candidate centromeric regions in ATCC 1015. Candidates were identified within the middle 60% of each chromosome. Coordinates are 1-based and inclusive, displayed in kb. Gene densities refer to the candidate (in) or whole chromosome (chr); insertion densities refer to the candidate (in) or the rest of the chromosome (out), with fold depletion calculated as out/in. Sites are distinct pooled junctions. All candidates contain 0.00% assembly Ns. Aligned percentages are the union of query intervals aligned to CBS 513.88 with minimap2 -x asm5. The final column gives the distance in ATCC 1015 coordinates from the nearest candidate boundary to the mapped midpoint of a sequence flanking the published CBS 513.88 centromere junction [22]. *n.p.*, no immediate flanking-anchor support: Chr3 lacked a qualifying anchor, and the retained Chr7 anchor lay 400 kb from the published junction.

| Chr | Region (kb) |  |  | Genes / 100 kb |  | Sites / kb |  |  | Aligned % |  | CBS 513.88<br>CBS ctr (kb) |
| --- | --- | --- | --- | --- | --- | --- | --- | --- | --- | --- | --- |
|  | start | end | GC % | in | chr | in | out | fold | chr | region |  |
| Chr1 | 1,610 | 1,703 | 17.2 | 0.0 | 32.5 | 7.2 | 66.6 | 9.3 | 96.3 | 4.2 | 10.6 |
| Chr2 | 1,102 | 1,189 | 17.2 | 0.0 | 31.5 | 5.9 | 80.5 | 13.7 | 94.3 | 6.6 | 13.1 |
| Chr3 | 1,978 | 2,064 | 16.9 | 0.0 | 31.0 | 5.2 | 73.5 | 14.1 | 95.6 | 2.0 | n.p. |
| Chr4 | 2,443 | 2,537 | 18.2 | 2.1 | 30.7 | 6.8 | 76.8 | 11.4 | 97.2 | 6.1 | 3.5 |
| Chr5 | 2,463 | 2,550 | 17.5 | 0.0 | 29.8 | 4.7 | 63.2 | 13.4 | 86.5 | 14.4 | 4.0 |
| Chr6 | 1,629 | 1,724 | 17.8 | 0.0 | 30.7 | 5.9 | 72.6 | 12.4 | 96.8 | 5.2 | 6.5 |
| Chr7 | 2,804 | 2,898 | 17.6 | 1.1 | 31.4 | 5.7 | 81.3 | 14.2 | 96.2 | 17.0 | n.p. |
| Chr8 | 1,604 | 1,694 | 17.5 | 0.0 | 32.0 | 5.3 | 77.8 | 14.7 | 97.8 | 2.1 | 9.0 |

## Appendix B Supplementary Table S2: Insertion density and chromatin accessibility

**Table S2 Supplementary Table S2.** Association between insertion density and published ATAC-seq signal. Spearman correlations compare pooled insertion junctions remapped to CBS 513.88 with published CBS 513.88 ATAC-seq cut sites [19], using 3 kb windows in the CBS 513.88 chromosome coordinate system. The pooled analysis includes all 11,335 windows. Confidence intervals use 2,000 block-bootstrap resamples of 33 consecutive windows. The rotation null uses 2,000 chromosome-wise circular shifts of the ATAC vector to break alignment between the signals while retaining their spatial structure. The null mean is a correlation coefficient, not a *P* value; plus-one *P* values are given below the table. The final columns restrict correlations to windows with both signals above zero. Random seed: 20260921. Analysis script:

| | Windows | $\rho$ | 95 % CI | | Null mean<br>$\rho$ | Both<br>> 0 | $\rho$ (both<br>> 0) |
| --- | --- | --- | --- | --- | --- | --- | --- |
| | $n$ | | low | high | | $n$ | |
| Pooled | 11,335 | 0.629 | 0.602 | 0.654 | 0.003 | 11,001 | 0.642 |
| Chr1 | 1,111 | 0.679 | 0.623 | 0.728 | < 0.001 | 1,084 | 0.685 |
| Chr2 | 1,512 | 0.655 | 0.599 | 0.707 | < 0.001 | 1,506 | 0.664 |
| Chr3 | 1,483 | 0.632 | 0.579 | 0.685 | -0.001 | 1,447 | 0.631 |
| Chr4 | 2,023 | 0.698 | 0.655 | 0.742 | < 0.001 | 2,015 | 0.701 |
| Chr5 | 998 | 0.682 | 0.611 | 0.743 | 0.001 | 976 | 0.694 |
| Chr6 | 1,379 | 0.548 | 0.469 | 0.627 | -0.002 | 1,300 | 0.563 |
| Chr7 | 1,110 | 0.606 | 0.533 | 0.679 | < 0.001 | 1,045 | 0.586 |
| Chr8 | 1,719 | 0.566 | 0.448 | 0.667 | < 0.001 | 1,628 | 0.622 |

Every row has $P < 0.01$ against the rotation null: the largest is 0.0015 on Chr1 and Chr7, and the pooled value is 0.0005, the resolution floor of 2,000 rotations. The null mean is $\rho$ under the rotations and is near zero throughout, providing a reference for the observed spatial correspondence.

## Appendix C Supplementary Table S3: Opposing stable classifications between carbon sources

**Table S3:** Genes with opposing stable classifications between glucose and xylose. All 91 reversals are included. G/X denote glucose/xylose; ESS/NESS denote stable predicted essential/nonessential classes. V2 means are classification scores across 1,000 fits in V2 repeated fitting, not measured fitness. Stable ESS requires mean*>* 0.8 and at least 90% of scores *>* 0.5; stable NESS requires mean *<* 0.2 and at least 90% of scores *<* 0.5. A dash indicates no accepted UniProt mapping.

| ATCC 1015 locus ID | UniProt | Function | G class | X class | G mean | X mean |
| --- | --- | --- | --- | --- | --- | --- |
| ANIGTU_000054 | — | hypothetical protein | ESS | NESS | 0.917 | 0.151 |
| ANIGTU_000310 | G3XMU2 | Chitin synthase | ESS | NESS | 0.815 | 0.118 |
| ANIGTU_000684 | G3XNU9 | Major facilitator superfamily (MFS) profile domain-containing protein | ESS | NESS | 0.803 | 0.072 |
| ANIGTU_000781 | — | hypothetical protein | ESS | NESS | 0.844 | 0.121 |
| ANIGTU_000916 | G3XN85 | hypothetical protein | ESS | NESS | 0.856 | 0.039 |
| ANIGTU_001470 | G3XPP0 | DNA repair protein rhp7 treble clef domain-containing protein (Fragment) | ESS | NESS | 0.807 | 0.142 |
| ANIGTU_001860 | G3XRW7 | ER membrane protein complex subunit 6 | ESS | NESS | 0.807 | 0.194 |
| ANIGTU_002034 | G3XSR9 | Mannan endo-1,6-alpha-mannosidase | ESS | NESS | 0.804 | 0.069 |
| ANIGTU_002252 | G3XTD1 | hypothetical protein | ESS | NESS | 0.824 | 0.120 |
| ANIGTU_002587 | — | hypothetical protein | ESS | NESS | 0.836 | 0.088 |
| ANIGTU_002686 | G3XS91 | Citrate exporter 1 (Fragment) | ESS | NESS | 0.824 | 0.064 |
| ANIGTU_002736 | — | hypothetical protein | ESS | NESS | 0.884 | 0.156 |
| ANIGTU_002972 | G3XW93 | ERCC4 domain-containing protein | ESS | NESS | 0.864 | 0.182 |
| ANIGTU_003215 | G3XVJ7 | Prohibitin | ESS | NESS | 0.914 | 0.154 |
| ANIGTU_003325 | — | C6 transcription factor | ESS | NESS | 0.903 | 0.149 |
| ANIGTU_003915 | G3XWZ6 | hypothetical protein | ESS | NESS | 0.861 | 0.172 |
| ANIGTU_003938 | G3XX20 | Alpha N-terminal protein methyltransferase 1 | ESS | NESS | 0.953 | 0.090 |
| ANIGTU_004093 | G3XZY9 | Intradiol ring-cleavage dioxygenases domain-containing protein | ESS | NESS | 0.934 | 0.178 |
| ANIGTU_004233 | — | hypothetical protein | ESS | NESS | 0.831 | 0.123 |
| ANIGTU_004518 | G3XZ09 | hypothetical protein | ESS | NESS | 0.821 | 0.071 |
| ANIGTU_004661 | — | Oxysterol-binding protein OBPa | ESS | NESS | 0.831 | 0.080 |
| ANIGTU_004929 | G3Y2N6 | histidine kinase | ESS | NESS | 0.827 | 0.088 |
| ANIGTU_004953 | G3Y2G5 | hypothetical protein | ESS | NESS | 0.911 | 0.195 |
| ANIGTU_004960 | G3Y2G1 | DUF1446 domain protein | ESS | NESS | 0.934 | 0.130 |
| ANIGTU_005573 | G3Y348 | C2H2-type domain-containing protein | ESS | NESS | 0.854 | 0.183 |
| ANIGTU_005751 | G3Y289 | 1,3-beta-glucanosyltransferase | ESS | NESS | 0.895 | 0.101 |
| ANIGTU_006112 | G3Y556 | VOC domain-containing protein | ESS | NESS | 0.863 | 0.167 |
| ANIGTU_006238 | — | hypothetical protein | ESS | NESS | 0.808 | 0.136 |
| ANIGTU_006400 | — | hypothetical protein | ESS | NESS | 0.817 | 0.098 |
| ANIGTU_006414 | G3Y608 | Vacuolar protein sorting-associated protein 35 | ESS | NESS | 0.866 | 0.160 |
| ANIGTU_006813 | G3Y4R2 | Peptidyl-tRNA hydrolase | ESS | NESS | 0.865 | 0.183 |
| ANIGTU_006819 | G3Y4Q1 | Histone-lysine N-methyltransferase, H3 lysine-36 specific | ESS | NESS | 0.807 | 0.180 |
| ANIGTU_008281 | — | CCA tRNA nucleotidyltransferase | ESS | NESS | 0.833 | 0.141 |
| ANIGTU_008284 | — | E3 ubiquitin-protein ligase rad18 | ESS | NESS | 0.848 | 0.069 |
| ANIGTU_008838 | G3YCY9 | Class I glutamine amidotransferase-like protein | ESS | NESS | 0.871 | 0.166 |
| ANIGTU_009017 | G3YAX0 | hypothetical protein | ESS | NESS | 0.918 | 0.139 |
| ANIGTU_009441 | — | Protein required for ethanol metabolism | ESS | NESS | 0.911 | 0.125 |
| ANIGTU_009564 | G3XUF0 | Malformin synthetase mlfA | ESS | NESS | 0.808 | 0.159 |
| ANIGTU_010316 | — | hypothetical protein | ESS | NESS | 0.892 | 0.109 |
| ANIGTU_011126 | G3YFN9 | Iron-sulfur clusters transporter ATM1, mitochondrial (Fragment) | ESS | NESS | 0.814 | 0.163 |
| ANIGTU_011223 | G3YG67 | CFEM domain-containing protein | ESS | NESS | 0.830 | 0.037 |
| ANIGTU_011238 | G3YGC7 | Translation initiation factor eIF2B subunit gamma | ESS | NESS | 0.831 | 0.081 |
| ANIGTU_000185 | G3XMH0 | Demethylmenaquinone methyltransferase family protein | NESS | ESS | 0.149 | 0.837 |
| ANIGTU_000415 | — | ubiquitin-specific protease ubp15 | NESS | ESS | 0.113 | 0.844 |
| ANIGTU_000499 | — | Non-classical phosphatidylinositol transfer protein (PITP) | NESS | ESS | 0.173 | 0.844 |
| ANIGTU_000737 | — | serine/threonine protein kinase | NESS | ESS | 0.178 | 0.805 |
| ANIGTU_000798 | G3XPF5 | DUF7703 domain-containing protein | NESS | ESS | 0.200 | 0.831 |
| ANIGTU_001503 | — | hypothetical protein | NESS | ESS | 0.161 | 0.808 |
| ANIGTU_001892 | — | DnaJ-domain containing protein | NESS | ESS | 0.178 | 0.881 |
| ANIGTU_002004 | G3XSN8 | DUF221 domain protein | NESS | ESS | 0.136 | 0.851 |
| ANIGTU_002244 | — | high-affinity iron permease | NESS | ESS | 0.076 | 0.842 |
| ANIGTU_002952 | G3XWA9 | Extracellular exo-inulinase inuE | NESS | ESS | 0.078 | 0.828 |
| ANIGTU_003322 | — | hypothetical protein | NESS | ESS | 0.148 | 0.850 |
| ANIGTU_003534 | G3XXP9 | GTPase-activating protein GYP7 | NESS | ESS | 0.191 | 0.815 |
| ANIGTU_003555 | G3XXR8 | Peptidase M20 domain-containing protein 2 (Fragment) | NESS | ESS | 0.103 | 0.826 |
| ANIGTU_003680 | G3XY40 | Kinesin family protein | NESS | ESS | 0.191 | 0.825 |
| ANIGTU_003970 | — | Ankyrin | NESS | ESS | 0.070 | 0.803 |
| ANIGTU_004771 | — | hypothetical protein | NESS | ESS | 0.162 | 0.837 |
| ANIGTU_004831 | G3Y377 | Ubiquitin-like modifier-activating enzyme ATG7 (Fragment) | NESS | ESS | 0.175 | 0.802 |
| ANIGTU_005104 | — | extracellular serine-rich protein | NESS | ESS | 0.081 | 0.872 |
| ANIGTU_005211 | — | hypothetical protein | NESS | ESS | 0.182 | 0.879 |
| ANIGTU_005221 | G3Y124 | NAD(P) transhydrogenase, mitochondrial (Fragment) | NESS | ESS | 0.088 | 0.870 |
| ANIGTU_005326 | G3Y0N7 | hypothetical protein | NESS | ESS | 0.177 | 0.873 |
| ANIGTU_005400 | G3Y3P2 | Gfo/Idh/MocA-like oxidoreductase N-terminal domain-containing protein | NESS | ESS | 0.155 | 0.831 |
| ANIGTU_005849 | — | Mitochondrial dicarboxylate transporter | NESS | ESS | 0.190 | 0.889 |
| ANIGTU_006155 | G3Y599 | HSF-type DNA-binding domain-containing protein | NESS | ESS | 0.169 | 0.913 |
| ANIGTU_006298 | G3Y5P2 | Monooxygenase | NESS | ESS | 0.141 | 0.886 |
| ANIGTU_006919 | — | THO2 plays a role in transcriptional elongation | NESS | ESS | 0.103 | 0.833 |
| ANIGTU_007016 | G3Y446 | hypothetical protein | NESS | ESS | 0.148 | 0.888 |
| ANIGTU_007212 | — | C6 transcription factor | NESS | ESS | 0.064 | 0.806 |
| ANIGTU_007548 | — | monooxygenase | NESS | ESS | 0.070 | 0.820 |
| ANIGTU_007790 | — | hypothetical protein | NESS | ESS | 0.068 | 0.862 |
| ANIGTU_007893 | G3Y8Y6 | WD repeat protein | NESS | ESS | 0.071 | 0.908 |
| ANIGTU_008121 | G3Y9J9 | DNA binding protein SART-1 | NESS | ESS | 0.171 | 0.825 |
| ANIGTU_008143 | — | hypothetical protein | NESS | ESS | 0.143 | 0.905 |
| ANIGTU_008317 | G3YAG0 | alpha-L-rhamnosidase | NESS | ESS | 0.097 | 0.819 |
| ANIGTU_008341 | — | Glycosyltransferase Family 2 protein | NESS | ESS | 0.188 | 0.881 |
| ANIGTU_008483 | G3YBN9 | Arrestin C-terminal-like domain-containing protein | NESS | ESS | 0.125 | 0.827 |
| ANIGTU_008580 | G3YC71 | RRM domain-containing protein | NESS | ESS | 0.167 | 0.831 |
| ANIGTU_008708 | G3YCK8 | Carboxylic ester hydrolase | NESS | ESS | 0.106 | 0.816 |
| ANIGTU_009061 | G3YB12 | S1/P1 Nuclease | NESS | ESS | 0.140 | 0.839 |
| ANIGTU_009651 | — | catalytic protein | NESS | ESS | 0.097 | 0.815 |
| ANIGTU_010230 | G3YEH4 | Chorismate synthase (Fragment) | NESS | ESS | 0.173 | 0.860 |
| ANIGTU_010271 | G3YEL4 | Proteasome component | NESS | ESS | 0.150 | 0.936 |
| ANIGTU_010354 | G3YFH5 | Dystroglycan-type cadherin-like domain-containing protein | NESS | ESS | 0.113 | 0.927 |
| ANIGTU_010389 | G3YFQ8 | hypothetical protein | NESS | ESS | 0.164 | 0.936 |
| ANIGTU_010461 | G3YG23 | Nop domain-containing protein | NESS | ESS | 0.123 | 0.881 |
| ANIGTU_010515 | — | hypothetical protein | NESS | ESS | 0.121 | 0.831 |
| ANIGTU_010541 | — | hypothetical protein | NESS | ESS | 0.075 | 0.846 |
| ANIGTU_010907 | — | UV-damaged DNA binding protein | NESS | ESS | 0.094 | 0.827 |
| ANIGTU_011253 | — | hypothetical protein | NESS | ESS | 0.048 | 0.906 |

## Appendix D Supplementary Methods: analysis definitions and parameter records

### D.1 Reconstruction, compartment assignments and assay media

The accompanying source package contains the executable analysis and its input tables. Paths below are relative to source/packages/. Reconstruction medium bounds are recorded in reconstruction/data/media/growth media.tsv; carbon-exchange identities and the seven tested carbon sources are specified in reconstruction/config/modeling.json. In the RAVEN precursor-demand and biomass checks, a feasible optimum above 10*^−^*^6^ constituted production, with mass-balance and bound residuals assessed against a 10*^−^*^7^ flux tolerance. These decision tolerances are distinct from the optimization bounds used during gap filling.

Compartment expansion used the union of predicted locations of proteins associated with each non-transport reaction. DeepLoc membrane-association predictions distinguished membranes from the corresponding aqueous compartments. Reactions without localization evidence and transport reactions retained their original placement; the pyruvate-dehydrogenase complex was retained as one mitochondrial matrix unit. Protein assignments and the explicit exception are recorded in reconstruction/data/localization/and reconstruction/workflow/step04 ijb1325/expand compartments.py.

The aqueous thermodynamic parameters were as follows. Concentration bounds apply in mol L*^−^*^1^, ionic strength in mol L*^−^*^1^, and relative electrical potential in mV. Temperature was 298.15 K. Reaction and metabolite energy uncertainties were retained in the serialized thermodynamic model rather than replaced by a single uncertainty for all compounds.

| Compartment | pH | Ionic strength | Concentration range | Potential |
| --- | --- | --- | --- | --- |
| Cytoplasm | 7.60 | 0.25 | $10^{-6}$–0.05 | 0 |
| Extracellular | 5.00 | 0.00 | $10^{-8}$–0.10 | 60 |
| Mitochondrion | 7.50 | 0.25 | $10^{-6}$–0.05 | –180 |
| Nucleus | 7.00 | 0.00 | $10^{-6}$–0.05 | –15 |
| Peroxisome | 8.20 | 0.00 | $10^{-6}$–0.05 | 0 |
| Endoplasmic reticulum | 7.20 | 0.00 | $10^{-6}$–0.05 | 0 |
| Golgi | 6.35 | 0.00 | $10^{-6}$–0.05 | 0 |
| Vacuole | 6.20 | 0.00 | $10^{-6}$–0.05 | 0 |

The six compartments excluded from aqueous thermodynamic constraints were the cell envelope, lipid particle, and endoplasmic-reticulum, Golgi, mitochondrial and vacuolar membranes. Parameters are recorded in constraint models/config/parameters. json. The glucose reference medium used for thermodynamic growth restoration is identified there as formal glucose. For the gene-essentiality assays, each model’s complete exchange bounds are supplied in fig5/source data/rerun/, within each configuration’s medium.tsv; these include the native reaction identifiers, lower and upper bounds, and carbon-source condition. The model metadata and reaction inventories in the same directories preserve the original objectives and gene associations.

### D.2 Insertion units, sequence mappability and chromatin comparison

A distinct insertion site was defined by chromosome, position and strand. Condition-level profiles combined the two corresponding libraries by set union; pooled profiles combined all six libraries by set union. Thus, repeated recovery of the same site did not add another site to a combined profile. The coding-sequence mappability screen used the fraction of coding bases whose 25-mer had a unique placement in the assembly, considering both strands. Genes below 95% callable coding bases were excluded from insertion-absence and differential-count screens. The common reference-based gene mask is supplied in fig4/data/k25 per gene callable.tsv, and its construction in fig4/scripts/build mappability.py. This criterion does not require observing an insertion and is separate from the nearby-insertion requirement used for the two shared insertion-absence groups.

For the ATAC comparison, insertion-junction reads and published ATAC-seq reads were mapped to the CBS 513.88 assembly (GCA 000230395.2), rather than directly correlating positions from different strain assemblies. Supercontigs were arranged in the published chromosome order. Counts of distinct insertion junctions and ATAC cut sites were compared in the resulting 11,335 chromosome windows of 3 kb. The main correlation included zero-count windows; a sensitivity analysis retained only windows with both signals above zero. Confidence intervals used 2,000 resamples of consecutive 33-window blocks within each chromosome. The spatial null used 2,000 independent chromosome-wise circular shifts of the ATAC track, retaining spatial structure within each signal while breaking correspondence between them. Source tracks and the analysis are supplied in fig3/source data/and fig3/scripts/14 atac spatial null.py.

### D.3 Classifier feature panels, score adjustment and reference labels

The reference-label table is classifier v2 1000/TRAINING LABELS 1340.tsv; it records the transferred ESS/NESS class, FungiDB identifier and ATCC 1015 locus. Condition-specific input matrices retain both replicates as separate feature columns, with shared restriction-sequence features included once. Feature names, reference-label indices and original labels are preserved in classifier v2 1000/CURRENT INPUT G. npz and its xylose counterpart. The executable feature-panel and score definitions are in classifier v2 1000/scripts/run R013 new.py.

The binary score-adjustment gate was calculated separately for glucose and xylose from all 11,113 annotated genes, without using reference labels. A reliable-site subset required, in both replicates, at least five coding insertion sites, an effective-site fraction of at least 0.35 and a maximum digest-specific read Gini coefficient no greater than 0.85. Threshold quantiles were calculated within this subset. Gate-positive genes additionally required the geometric-mean reads-per-site measure to be at or below its 25th percentile, each replicate’s reads-per-site measure at or below its median, and geometric-mean site count and central-coding-region density at or below their respective 80th percentiles. The geometric-mean effective-site fraction and observed-to-expected site ratio had to be at or above their medians, while the maximum read Gini coefficient had to be at or below its median. The reads-per-site measure was log(1 +reads) *−* log(1 +sites) per replicate, averaged across replicates; the observed-to-expected ratio used the geometric-mean observed count divided by the geometric-mean locally expected count plus 0.5. The exact feature names and gate implementation are supplied in classifier v2 1000/scripts/historical gate.py.

V1 score classes determined retention, not replacement of the transferred reference labels. For example, on glucose V1 assigned 513 genes to the high-rank class, 640 to the low-rank class and 187 to the inconclusive class. The 1,153 retained genes nevertheless carried 504 ESS and 649 NESS original reference labels in V2. This distinction explains the different V1 score-class and V2 reference-label counts in Fig. 4E–F. The frozen retention masks are supplied as V1 KEEP G.npy and V1 KEEP X.npy in the classifier package.

